# Rearing conditions bidirectionally modulate cognitive abilities and AP-1 signaling in hippocampal neurons in a cell type-specific manner

**DOI:** 10.1101/2025.02.11.637399

**Authors:** Marta Alaiz-Noya, Federico Miozzo, Miguel Fuentes-Ramos, Magdalena Machnicka, Marcelina Kurowska, Macarena L. Herrera, Beatriz del Blanco, Sergio Ninerola, Isabel Bustos-Martínez, Bartek Wilczynski, Angel Barco

## Abstract

Environmental conditions profoundly influence cognitive development, particularly during early life. Transcriptional and epigenetic mechanisms may serve as molecular substrates for the lasting effects of environmental enrichment (EE) and impoverishment (IE) on cognitive abilities and hippocampal function. However, the specific gene programs driving these changes remain largely unknown. In this study, EE and IE modulated the cognitive abilities of mice in opposing directions. By combining hippocampal microdissection and genetic tagging of neuronal nuclei with genome-wide analyses of gene expression, chromatin accessibility, histone acetylation, and DNA methylation, we uncovered profound differences in the transcriptional and epigenetic profiles of CA1 pyramidal neurons and dentate gyrus (DG) granule neurons. These analyses revealed cell type-specific genomic changes induced by EE and IE, highlighting distinct patterns of neuroadaptation within each population. This multiomic screen pinpointed the activity-regulated transcription factor AP-1 as a crucial mediator of neuroadaptation to conditions during early life in both cell types, albeit through distinct downstream mechanisms. Conditional deletion of *Fos*, a core AP-1 subunit, in excitatory neurons hampered EE-induced cognitive enhancement, further underscoring the pivotal role of this transcription factor in neuroadaptation.

## Introduction

Cognitive development is profoundly influenced by rearing conditions. For instance, after Donald O. Hebb’s early discovery that domestic rats performed better in problem-solving tests than rats raised in laboratory cages (*1*), numerous studies have employed environmental enrichment (EE) to study the impact of a stimulating setting on cognitive development (*2*). Prolonged exposure to EE, which typically includes opportunity for physical exercise, exploration of novel objects and increased social interactions, has been shown to improve memory in various hippocampal-dependent tasks both in wild type animals and in a range of models of neuronal dysfunction, including brain injury, Huntington’s Disease, Alzheimer’s Disease and intellectual disability disorders (*3–8*). Investigation of the underlying cellular and synaptic mechanisms revealed that EE stimulates neurogenesis (*9–11*) and increases spine density, dendritic length and dendritic complexity in hippocampus and cortex regions (*3, 7, 12*). Conversely, an impoverished environment (IE), conceived as individual housing with no exposure to stimulating objects, has been used to model psychiatric disorders in rodents (*13–15*), and leads to cognitive impairments, decreased dendritic length and spine density, and defects in glutamatergic transmission (*16–19*).

The application of EE as a potential intervention for a broad range of human traits (*20, 21*), from neurodevelopmental disorders in children (*22*) to cognitive decline in the elderly (*23*), has sparked significant interest in understanding the molecular mechanisms by which housing conditions influence cognition. Transcriptional and epigenetic processes, which are well-established mediators between environmental factors and the genome (*24, 25*), play a crucial role in synaptic plasticity underlying learning and memory (*26–29*). These mechanisms are therefore appealing candidates for translating the impact of rearing conditions into cognitive outcomes. However, genome-wide studies on cortical and hippocampal neurons following EE have identified relatively few consistent changes in gene expression (*30–33*). The molecular study of IE has received far less attention, leaving the genomic modifications induced by IE largely unexplored. A comprehensive understanding of the epigenetic and transcriptional programs driving environment-induced changes in neuronal plasticity and behavior could be crucial for developing new therapeutic strategies that mimic EE or correct IE-related deficits. Yet, the underlying mechanisms remains elusive. Since cellular heterogeneity in nervous tissue may obscure subtle changes induced by environmental conditions, strategies that mitigate cellular heterogeneity would significantly enhance the sensitivity and specificity of epigenomic and transcriptional analyses, offering unprecedented insights.

In this study, we exposed female mice to EE and IE during the juvenile period to investigate how rearing conditions influence cognitive abilities and to map the associated transcriptional and chromatin changes in the two main hippocampal neuron types: CA1 pyramidal and DG granule neurons. Our behavioral experiments revealed opposite and durable effects on the animals’ cognitive abilities, while the multiomic analysis identified activity-regulated AP-1 as a central player, activated by EE and inhibited by IE, leading to the cell type-specific modulation of synaptic target genes critical for memory functions.

## Results

### Bidirectional regulation of cognitive capacities by rearing conditions

To investigate the impact of EE in cognitive functions, we developed a paradigm in which a large cohort of 3-week-old C57BL6/J female mice were weaned and housed together in an ample space with opportunities for voluntary exercise, social interaction and exploration of novel objects, where toys and running wheels were changed every two weeks. In parallel, to investigate the impact of social isolation and IE, 3-week-old female mice were weaned and housed individually in small cages, shielded from acoustic, visual, and social stimuli. After 3 months, we compared the performance of the EE and IE cohorts and their littermates housed in standard cages (SC; 4-5 mice per cage) (**Fig. 1A**).

**Figure 1.**
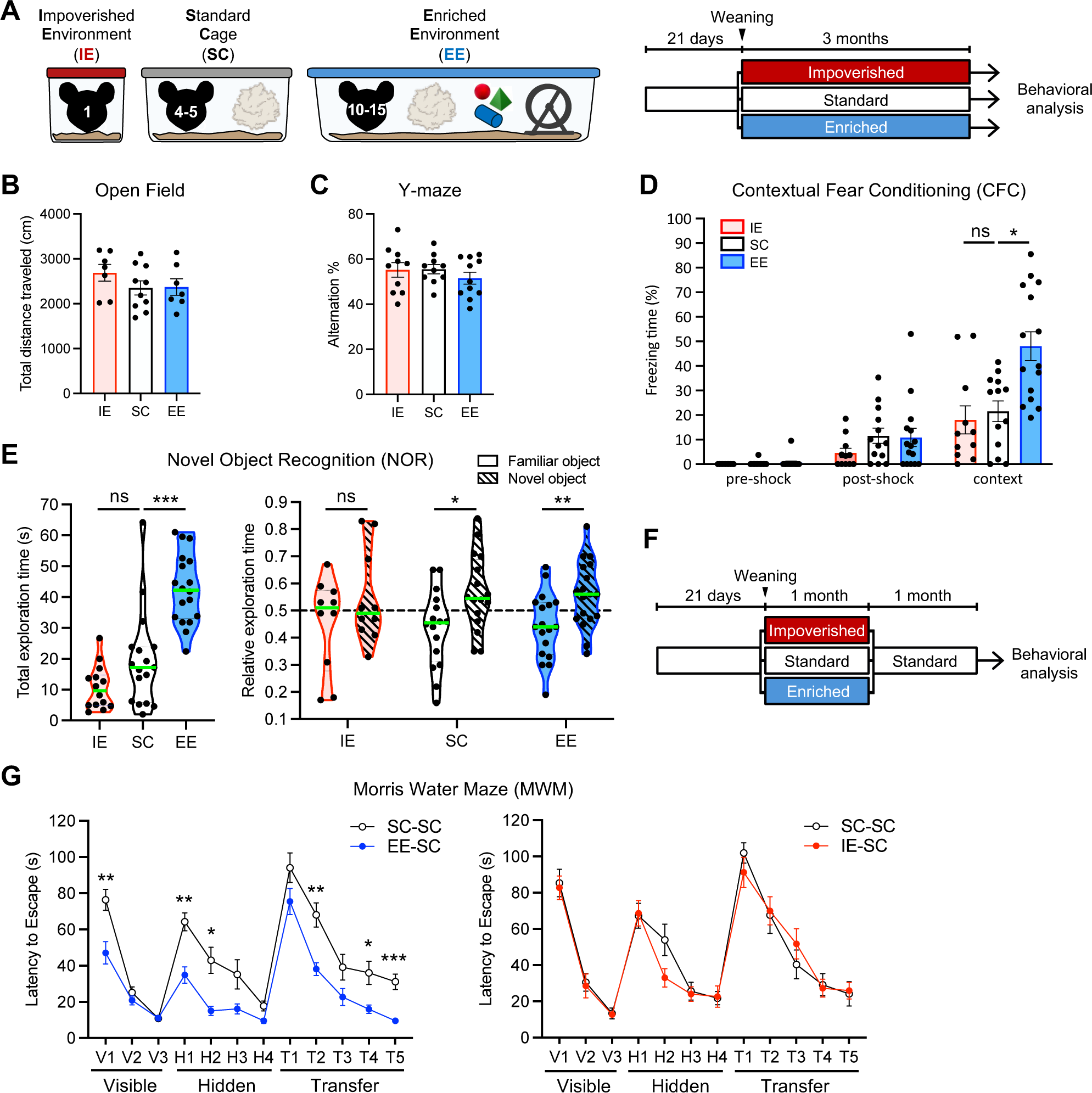
Bidirectional impact of rearing conditions on cognitive capacities. **A**. Experimental design. At weaning (P21), littermate WT female mice were randomly distributed in impoverished (IE, red), standard (SC, white), or enriched (EE, blue) environment for three months before behavioral analysis. **B**. Total distance traveled in the Open Field test. Mean ± SEM is shown. n = 7-10 mice per group. No statistically significant differences comparing EE vs SC and IE vs SC by Kruskal-Wallis test followed by Dunn’s multiple comparisons test. **C**. Alternation percentage in the Y-maze. Mean ± SEM is shown. n = 10-11 mice per group. No statistically significant differences comparing EE vs SC and IE vs SC by Kruskal-Wallis test followed by Dunn’s multiple comparisons test. **D**. Contextual Fear Conditioning. The freezing time before the shock, after the shock, and during the context session was measured. Mean ± SEM is shown. n = 11-15 mice per group. Kruskal-Wallis test followed by Dunn’s multiple comparisons test comparing EE vs SC and IE vs SC, * p < 0.05, ns not significant. **E**. Novel Object Recognition. The total time spent exploring the identical objects during the training (left) and the relative time spent exploring the familiar and the novel object during the test session (right) are shown. The green horizontal line across the violin plot represents the median. n = 14-18 mice per group. Mice with a total exploration time inferior to 5 s during the training or the test were excluded from the analysis of the relative exploration time. Total exploration time, Kruskal-Wallis test followed by Dunn’s multiple comparisons test comparing EE vs SC and IE vs SC, *** p < 0.001, ns not significant. Relative exploration time, Two-tailed Mann-Whitney test, * p < 0.05, ** p < 0.01, ns not significant. **F**. Experimental design. At weaning (P21), littermate female mice were randomly distributed in impoverished (IE-SC, red), standard (SC-SC, white), or enriched (EE-SC, blue) environment for one month, and were then kept in standard environment for one additional month before behavioral analysis. **G**. Morris Water Maze. The latency to reach the platform as an indicator of spatial memory was measured for EE-SC vs SC-SC (left) and IE-SC vs SC-SC (right). n = 10-14 mice per group. Repeated measures ANOVA with correction for multiple comparisons using Šídák’s test. EE-SC vs SC-SC, visible phase: day ****, rearing conditions ***. Hidden and reversal phase: day ****, rearing conditions ****. IE-SC vs SC-SC, visible, hidden and reversal phase: day ****, rearing conditions ns. * p < 0.05, ** p < 0.01, *** p < 0.001, **** p < 0.0001, ns not significant.

First, we examined mice basal exploratory activity in an open field test and did not find any significant change either in the total distance travelled, nor in the time spent in the center and the periphery of the arena (**Fig. 1B** and **Supp. Fig. S1A**). Exploration of a Y-maze did not reveal significant differences either (**Fig. 1C** and **Supp. Fig. S1B**). Long-term memory was analyzed in a contextual fear conditioning (CFC) task in which the mice received a single foot-shock. EE mice froze significantly longer during recall, indicating better contextual memory in line with previous studies (*6*), while IE mice behaved similarly to SC mice (**Fig. 1D**). Furthermore, training in a Novel Object Recognition (NOR) task revealed that EE mice spent more time exploring the objects, while IE mice displayed a trend towards reduced exploration (**Fig. 1E, left**). Both SC and EE mice dedicated more time to explore the novel object in the test session, while IE mice could not discriminate between the familiar and novel object, suggesting impaired recognition memory (**Fig. 1E, right**, and **Supp. Fig. S1C**).

To evaluate the persistence of cognitive changes induced by rearing conditions, we examined the performance of EE and IE mice after being returned to SC one month prior to behavioral assessment (henceforth referred to as EE-SC, IE-SC and SC-SC) (**Fig. 1F**). Given that spatial navigation is known to improve following EE (*5, 7, 34*), we first assessed the mice using the Morris Water Maze (WMW) test. EE-SC mice demonstrated superior learning compared to the SC-SC group, whereas IE-SC mice did not differ significantly from their controls (**Fig. 1G** and **Supp. Fig. S1D**). In the CFC task, EE-SC mice also exhibited significantly stronger associative memory compared to their SC-SC littermates, while IE-SC mice performed similarly to the SC-SC group (**Supp. Fig. S1E**). Similarly, in a working memory task conducted using the radial maze, EE-SC mice outperformed their SC-SC littermates, while IE-SC mice showed a greater number of errors (**Supp. Fig. S1F**).

In summary, our findings reveal that experiencing IE, and especially EE, during the juvenile period exerts a lasting impact on hippocampal-dependent memory processes, highlighting the potential existence of enduring molecular changes in hippocampal neurons.

### CA1 and DG excitatory neurons exhibit distinct transcriptional and chromatin profiles

The chromatin of CA1 pyramidal neurons and DG granule neurons, the two main neuronal populations in the hippocampus with fundamental yet distinct roles in memory formation (*35, 36*), are likely depository for environment-induced molecular changes that influences neuronal responsiveness and hippocampal circuits functioning. To investigate if EE and IE trigger changes in the transcriptome and chromatin of these neurons, we generated mice in which the nuclear envelope of forebrain excitatory neurons is fluorescently tagged with the fusion protein SUN1-GFP upon tamoxifen (TMX) injection (**Fig. 2A**; hereafter referred to as *Sun1-GFP* mice). This labeling enabled the efficient isolation of these neuronal nuclei using fluorescence activated nuclear sorting (FANS) for subsequent transcriptional and epigenomic analyses (*29*).

**Figure 2.**
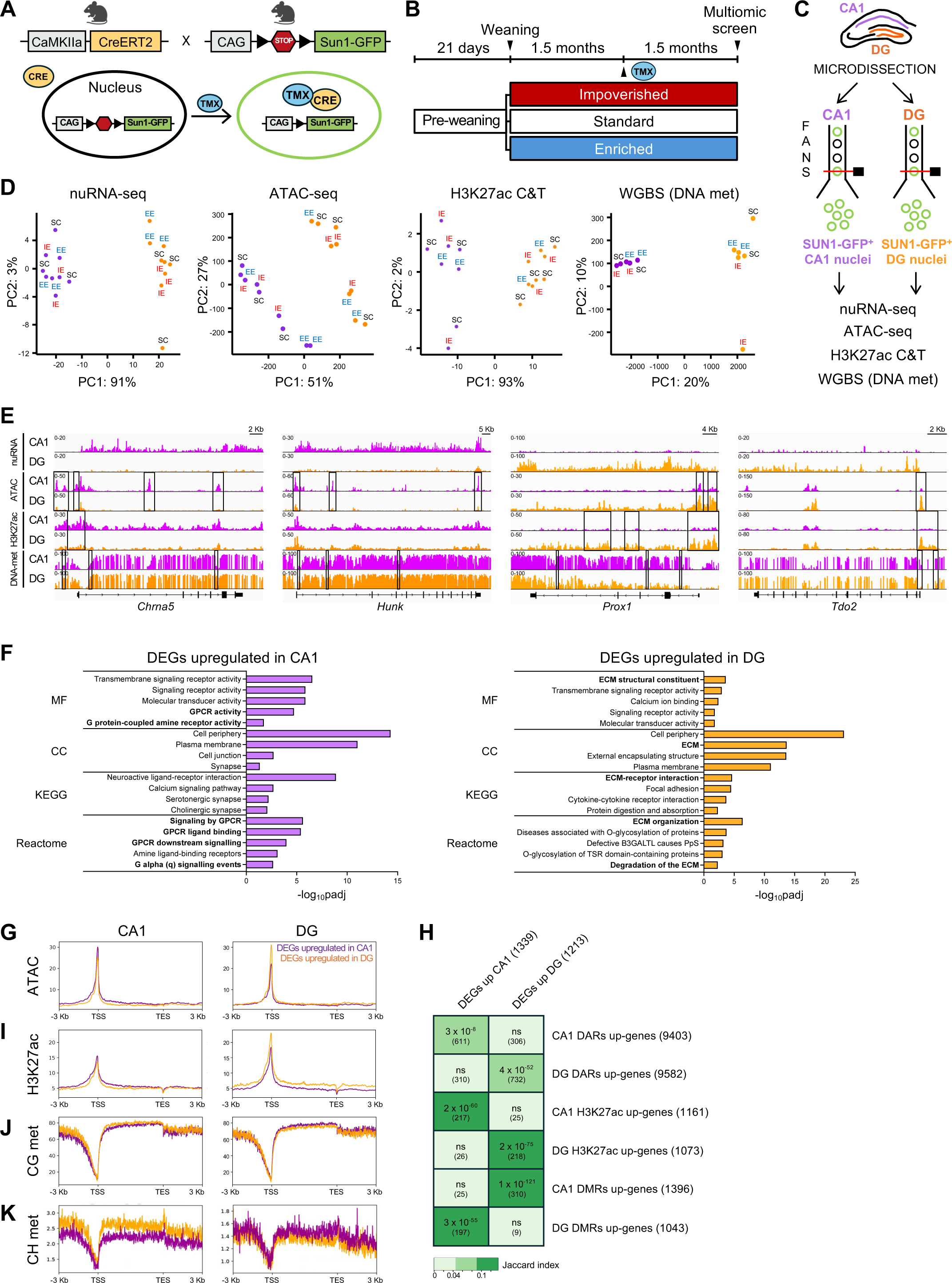
Multiomic analysis reveals profound and correlated differences in transcriptional and epigenetic profiles between CA1 pyramidal neurons and DG granule neurons. **A**. Genetic strategy for the conditional expression of *Sun1-GFP* in forebrain excitatory neurons. **B**. At weaning (P21), *Sun1-GFP* female mice were randomly distributed in enriched (EE, red), standard (SC, white) or impoverished (IE, red) environment for 3 months. At 1.5 month, mice were administered TMX. **C**. Experimental design. SUN1-GFP^+^ nuclei were isolated by FANS from manually dissected CA1 and DG regions for downstream multiomic analysis. **D.** PCA of nuRNA-seq, ATAC-seq, H3K27ac CUT&TAG and WGBS DNA methylation profiles of CA1 and DG SUN1-GFP^+^ neurons. n = 3 (nuRNA-seq, ATAC-seq), n = 2-4 (H3K27ac CUT&TAG) and n = 2 (WGBS DNA methylation) samples per region per rearing condition. **E**. Snapshot of nuRNA-seq, ATAC-seq, H3K27ac and DNA methylation genomic profiles of *Chrna5, Hunk* (CA1 pyramidal neurons markers), *Prox1* and *Tdo2* (DG granule neurons marker) genes in CA1 (violet) and DG (orange) samples from SC mice. DNA methylation is represented as the percentage of methylation at a given position. DARs, differential H3K27ac peaks and DMRs are highlighted by rectangles. The merged profiles of multiple independent samples are shown. **F**. Gene Ontology (GO) enrichment analysis of the up-regulated DEGs in CA1 (left) and DG (right) excitatory neurons. The 4-5 significant GO terms with higher -log_10_padj for Molecular Function (MF), Cellular Component (CC), KEGG pathway and Reactome pathway are shown. GO terms related to GPCR signaling and ECM are highlighted in bold for CA1 and DG, respectively. **G**. Density distribution of ATAC reads across the DEGs up-regulated in CA1 and DG. **H**. Matrix illustrating the overlap between the genes significantly up-regulated in CA1 or DG neurons, and the genes associated with DARs, H3K27ac peaks and DMRs significantly increased in CA1 or DG neurons. The Jaccard index and the number of shared genes for each comparison, in brackets, are indicated. ns, not significant. **I**. Density distribution of H3K27ac reads across the DEGs up-regulated in CA1 and DG. **J, K**. Percentage of cytosine methylation within CG context (J) and CH context (K) across the DEGs up-regulated in CA1 and DG.

*Sun1-GFP* mice were housed in IE, SC or EE starting at 3-weeks of age, and after 6 weeks, TMX was administered to trigger SUN1-GFP expression (**Fig. 2B**). After 6 additional weeks in their respective environments, the mice were euthanized, CA1 and DG hippocampal layers were microdissected, and the tissues dissociated and subjected to FANS for multiomic analysis (**Fig. 2C**). RT-qPCR assays of specific markers of DG (*Tdo2, Dsp*) and CA1 (*Ccn3*) (*37*) confirmed the efficiency and precision of the hippocampal subregions microdissection (**Supp. Fig. S2A**).

Transcriptional changes were assessed using nuRNA-seq, providing insights into gene expression dynamics. Chromatin accessibility was evaluated through ATAC-seq (assay for transposase-accessible chromatin coupled to sequencing), which allowed for the identification of regulatory elements and potential changes in the epigenetic landscape. Histone H3 lysine 27 acetylation (H3K27ac) was investigated using CUT&TAG (*38*), a targeted approach to detect modifications associated with active enhancers and promoters, linking chromatin state to transcriptional activity. DNA methylation patterns were examined via low-input whole-genome bisulfite sequencing (WGBS) to further identify changes in the epigenetic state influencing gene regulation. This multiomic strategy provided a robust framework for unraveling the interplay between housing conditions, gene expression and chromatin modifications. Principal component analysis (PCA) revealed a clear separation between CA1 (purple) and DG (orange) samples in all four genomic screens, while we did not observe any obvious clustering of the samples according to environmental conditions (**Fig. 2D**).

As a first step in our analyses, we compared CA1 and DG neurons in SC conditions to identify fundamental differences in their transcriptomic and chromatin profiles, before assessing how environmental perturbations modify these cell-type specific programs. nuRNA-seq identified 2,552 differentially expressed genes (DEGs; padj < 0.05 and log_2_FC(CA1/DG) > 1), with 1,339 genes up-regulated in CA1 and 1,213 genes up-regulated in DG (**Supp. Fig. S2B** and **Supp. Table 1**). DEGs included well-known markers of CA1 pyramidal neurons, such as *Chrna5* and *Hunk*, and DG granule neurons, such as *Prox1* and *Tdo2* (**Fig. 2E**). Gene ontology (GO) analysis revealed profound differences between the two neuronal types in terms of cell adhesion molecules and intracellular signaling. In particular, multiple G-protein-coupled (*Gprc6A, Gpr101, Gpr139, Gpr26, Gpr161, Gpr3*), serotoninergic (*Htr1b, Htr2c, Htr5b, Htr1f*), and cholinergic (*Chrm2, Chrm3, Chrm5, Chrna4*) receptors were found up-regulated in CA1 neurons, while extracellular matrix (ECM) proteins were highly overrepresented in DG neurons, including members of laminin (*Lama1, Lama2, Lama5*), collagen (*Col1a1, Col1a2, Col3a1, Col4a5, Col4a6, Col5a3, Col12a1, Col13a1, Col15a1, Col27a1*) and ADAMTS (short for *a disintegrin and metalloproteinase with thrombospondin motifs*) families (*Adamts3, Adamts12, Adamts15, Adamts17, Adamts18, Adamtsl2*) (**Fig. 2F** and **Supp. Table 2**). These broad transcriptomic differences likely underlie the distinct synaptic features and functional specialization of CA1 and DG excitatory neurons (*35, 39*).

ATAC-seq identified 39,084 differentially accessible regions (DARs; padj < 0.01 and log_2_FC(CA1/DG) > 1) when comparing CA1 and DG samples in the SC condition, with 20,465 regions more accessible in CA1 and 18,619 regions more accessible in DG (**Supp. Fig. S2C** and **Supp. Table 3**). Chromatin opening is a well-stablished feature of actively transcribed genes. In agreement, CA1 pyramidal neurons displayed higher accessibility across CA1 up-regulated genes than DG up-regulated genes, and *vice versa* for DG granule neurons (**Fig. 2E, G**). Furthermore, the genes annotated to DARs augmented in a specific cell-type showed a highly significant overlap with the DEGs upregulated in the same neuronal type (**Fig. 2H**).

When comparing CUT&TAG H3K27ac profiles between CA1 and DG samples, 2,426 and 1,765 peaks were retrieved as significantly increased in CA1 or DG neurons, respectively (padj < 0.05 and log_2_FC(CA1/DG) > 1) (**Supp. Fig. S2D** and **Supp. Table 4)**. Like chromatin accessibility, H3K27ac levels at CA1 up-regulated genes were higher than DG up-regulated genes in CA1 pyramidal neurons, and *vice versa* for DG granule neurons (**Fig. 2E, I**). Coherently, the genes annotated to H3K27ac peaks that increased in one cell-type largely overlapped with the DEGs upregulated in the same neuronal population (**Fig. 2H**). Overall, we found widespread changes in chromatin accessibility and H3K27 acetylation between pyramidal and granule neurons, which positively correlate with their cell-type specific transcriptional signatures, consistent with the established association of these chromatin features with active gene expression.

DNA methylation analysis using WGBS revealed even more numerous differences between the two cell types, identifying 91,987 and 82,159 differentially methylated regions (DMRs; FDR < 0.01 and methylation difference > 25 %) with higher methylation levels in CA1 and DG neurons, respectively (**Supp. Fig. S2E** and **Supp. Table 5)**. Promoter DNA methylation is typically associated with gene repression, whereas the effects of gene body methylation on transcription are more complex (*40*). In dividing cells, gene body DNA methylation correlates with gene expression; however, in non-dividing cells such as neurons, it does not correlate with active transcription (*41, 42*). We found that genes upregulated in CA1 exhibited regions with lower CG and CH methylation level in the chromatin of CA1 pyramidal neurons across both the promoter and gene body compared to genes upregulated in DG. Conversely, in DG granule neurons, lower DNA methylation levels were observed in the gene bodies of DG-upregulated genes (**Fig. 2E, J, K**). Consistent with this, and in contrast to genes annotated to ATAC and H3K27ac peaks, the genes associated with DMRs that augmented in one neuronal type significantly overlapped with DEGs upregulated in the other cell type (**Fig. 2H**). Taken together, our data show that CA1 and DG excitatory neurons exhibit extensive differences in DNA methylation profiles, which inversely correlate with their unique transcriptional programs, indicating a repressive role for this chromatin mark in regulating neuron type-specific gene expression.

To better understand how the unique chromatin profiles of CA1 and DG excitatory neurons result into cell-type specific transcriptional programs, we classified all the accessible regions identified by ATAC-seq in the two cell types into promoters or enhancers, based on the presence or absence of the epigenetic mark histone 3 lysine 4 trimethylation (H3K4me3), respectively. Our analysis determined 20,606 promoters and 138,191 enhancers in CA1 pyramidal neurons, and 19,039 promoters and 103,699 enhancers in DG granule neurons (**Supp. Fig. S3A**). Remarkably, a substantial fraction of promoters and enhancers were exclusive to one cell-type or the other, with 40% of pyramidal neuron enhancers (over 55,000 regions) which were absent in granule neurons, denoting extensive differences in chromatin regulatory landscapes between the two populations (**Supp. Fig. S3B**). The genes associated to these cell-type exclusive promoters and enhancers largely overlapped with DEGs upregulated in the respective neuronal populations, suggesting that they play a crucial role in shaping the distinct transcriptional profiles of CA1 and DG neurons (**Supp. Fig. S3C**). Additionally, DMRs with increased DNA methylation in one cell type were enriched in enhancers exclusive to the other, further supporting the repressive role of DNA methylation (**Supp. Fig. S3D**).

Altogether, our genomic analyses revealed profound differences in the transcriptional and chromatin profiles of the two main neuronal populations in the hippocampus. Furthermore, the strong correlation between transcriptomic and chromatin profiles within each cell population demonstrates the robustness and cell-type specificity of our datasets.

### Intrinsic differences in activity-driven transcription between pyramidal and granule neurons

To understand how TFs operate differently in CA1 and DG regulatory regions, we performed a TF footprint analysis of our ATAC-seq dataset within promoters and enhancers shared by CA1 and DG excitatory neurons (p < 0.05 and binding differential score > 0.25 or < -0.25). The analysis revealed that the activity-dependent transcriptional complexes AP-1 and NEUROD (*43, 44*) exhibited stronger predicted binding to chromatin in pyramidal neurons compared to granule neurons, both within promoters and enhancers (**Fig. 3A, B** and **Supp. Fig. S4A**). These results suggest that CA1 neurons exhibit distinct responses to synaptic activation compared to DG neurons, with activity-induced signaling pathways being more strongly activated in CA1 neurons under standard conditions, likely reflecting higher baseline activity in pyramidal neurons.

**Figure 3.**
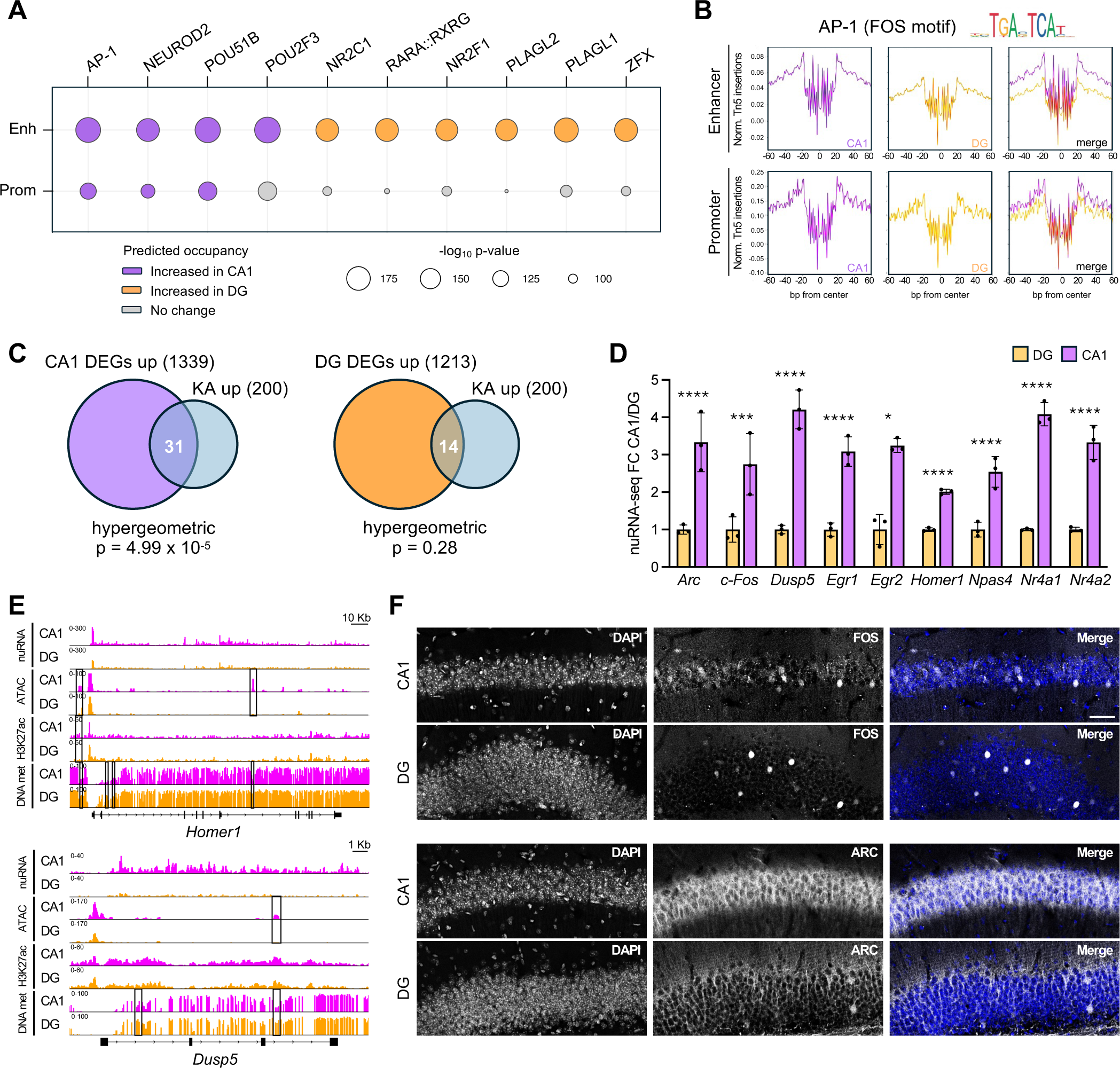
CA1 pyramidal neurons exhibit higher expression of the activity-induced transcriptional program than DG granule neurons. **A**. TF binding prediction in enhancers and promoters comparing CA1 (violet) and DG (orange) excitatory neurons in SC conditions (p < 0.05 and binding differential score > 0.25 or < -0.25). Circle size indicates motif enrichment p-value, and circle color refers to the predicted occupancy by the TFs. **B**. Digital footprint for AP-1 (FOS motif) in enhancers (top) and promoters (bottom) comparing CA1 and DG excitatory neurons. Values correspond to normalized Tn5 insertions. **C**. Venn diagrams showing the overlap between the DEGs up-regulated in CA1 (left) or DG (right) excitatory neurons and the 200 most up-regulated genes in response to KA treatment (*29*). **D**. Relative expression levels of a panel of IEGs in CA1 and DG excitatory neurons in SC conditions, as measured by nuRNA-seq. Data are expressed as fold change over DG neurons levels. n = 3 samples per region. Two-tailed Mann-Whitney test, * padj < 0.05, *** padj < 0.001, **** padj < 0.0001. **E**. Snapshot of nuRNA-seq, ATAC-seq, H3K27ac and DNA methylation genomic profiles of *Homer1* (top) and *Dusp5* (bottom) genes in CA1 (violet) and DG (orange) samples from SC mice. DNA methylation is represented as the percentage of methylation at a given position. DARs, differential H3K27ac peaks and DMRs are highlighted by rectangles. The merged profiles of multiple independent samples are shown. **F**. Representative immunostaining of FOS (top) and ARC (bottom) proteins in CA1 and DG coronal sections of SC mice. DNA was counterstained with DAPI. Scale bar, 40 µm.

To further investigate the regulation of activity-induced gene expression in the two cell types, we examined the levels of previously identified activity-induced genes (*29*) within our datasets. Interestingly, a significant fraction of the top 200 induced genes was upregulated in CA1 neurons compared to DG neurons in SC (**Fig. 3C**), including classical IEGs such as *Arc, Fos, Dusp5, Egr1, Egr2, Homer1, Npas4, Nr4a1* and *Nr4a2* (**Fig. 3D**). Some of these genes also displayed higher accessibility, increased H3K27ac levels and reduced DNA methylation in CA1 compared to DG (e.g., *Homer1* and *Dusp5*, **Fig. 3E**). Since IEGs are induced in response to neuronal activity, our results suggest that a larger fraction of CA1 neurons are active in standard housing conditions compared to DG neurons.

Consistently with the multiomic analysis, immunostaining of hippocampal sections revealed a moderate but spread expression of FOS protein in CA1 pyramidal neurons, while only a few sparse granule neurons in the DG layer displayed detectable FOS levels (**Fig. 3F, top**). Similarly, ARC showed a stronger expression in CA1 compared to DG (**Fig. 3F, bottom**).

These findings align with differences observed during flow cytometry. While DG nuclei showed a narrow green fluorescence distribution across all conditions, CA1 SC samples displayed a bimodal signal distribution, which was shifted toward higher levels after EE (**Supp. Fig. S5A, B**). *Sun1-GFP* expression is driven by the chimeric CAG promoter that contains binding sites for activity-regulated transcription factors (TFs) like the cAMP response element binding protein (CREB) and the activator protein 1 (AP-1) (*45*). Consistently, *Sun1-GFP* transcripts were significantly increased in animals treated with the glutamate receptor agonist kainic acid (KA) compared to saline (*29*) (**Supp. Fig. S5C**). Altogether, these results suggest that CA1 pyramidal neurons have a more sustained baseline activity than DG granule neurons, and that EE causes an increase in tonic activity-dependent transcription in CA1 neurons.

### EE enhances and IE downregulates the activity-induced transcriptional program in hippocampal neurons

Next, we conducted a detailed analysis of the transcriptional and chromatin differences induced by rearing conditions in the two cell types. In the PCA of CA1 nuRNA-seq samples, the experimental groups did not cluster distinctly, although IE nuclei tended to separate from the other two conditions (**Fig. 4A**). Indeed, IE was the primary driver of the transcriptional changes, leading to the downregulation of most DEGs (**Fig. 4B**). Notably, nearly all DEGs were more strongly expressed in CA1 than DG in standard conditions. We identified 58 DEGs (padj < 0.1; **Supp. Table 6**) which displayed a highly significant number of physical and functional interactions among them, including a group of 6 highly interconnected IEGs involved in transcriptional regulation (**Supp. Fig. S6A**). Remarkably, we noticed that numerous IEGs (*Arc, Fos, Fosb, Egr1, Egr2, Npas4, 1700016p03rik*) clustered together and were downregulated after IE and upregulated in response to EE (**Fig. 4B, C**). Consistent with flow cytometry findings, this evidence indicates that rearing conditions bidirectionally regulate the activity-dependent transcriptional program in CA1 pyramidal neurons.

**Figure 4.**
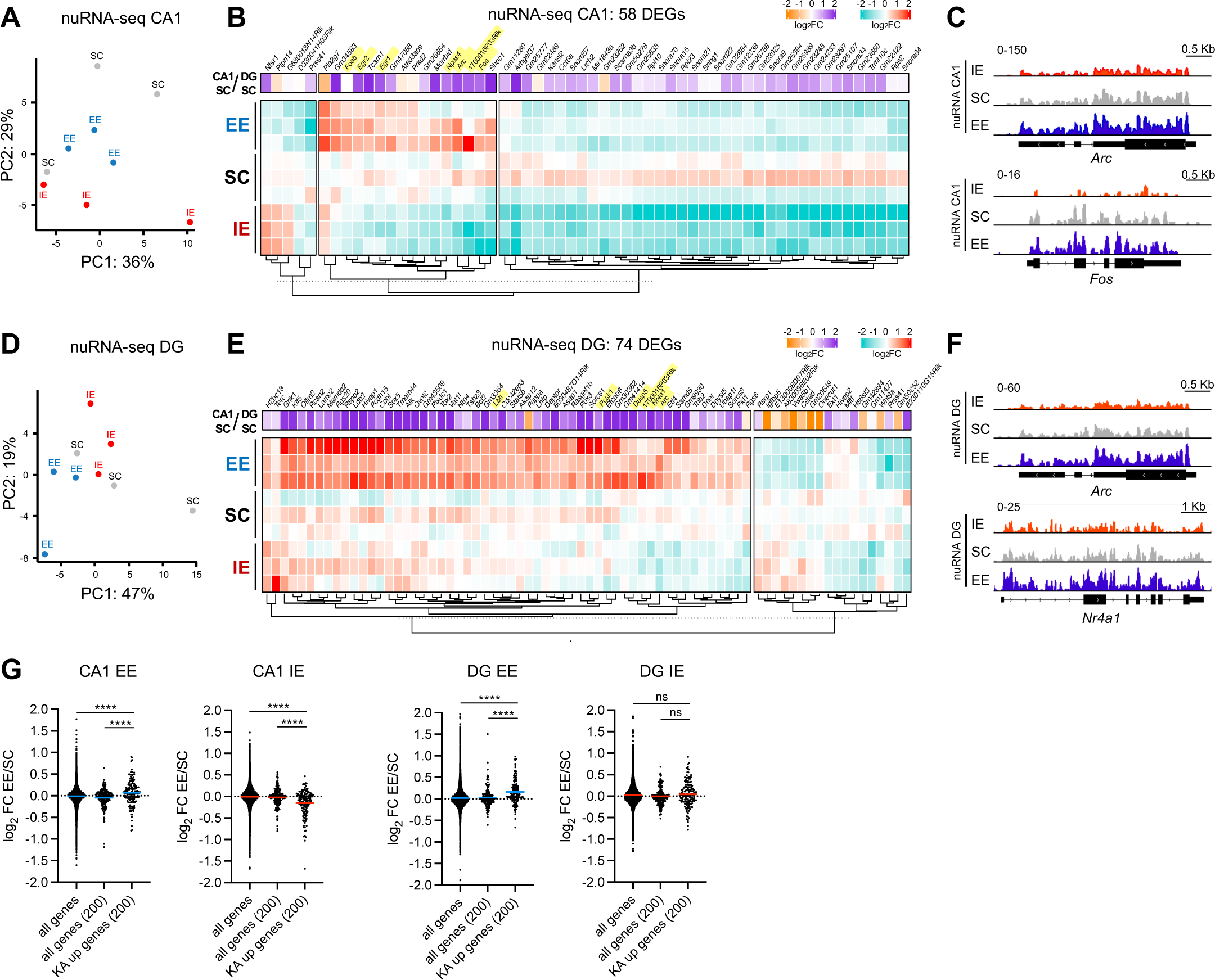
EE and IE modulate the activity-induced transcriptional program in CA1 and DG excitatory neurons in a cell-type specific manner. **A**. PCA of nuRNA-seq profiles of CA1 neurons in IE, SC an EE conditions (n =3 samples per rearing condition). **B**. Heatmap of DEGs in nuRNA-seq profiles of CA1 neurons in IE, SC and EE conditions (padj < 0.1). The gene expression fold change of each gene in the CA1 SC vs DG SC comparison (Fig. 2) is also shown. IEGs are highlighted in yellow. **C**. Snapshot of nuRNA-seq genomic profiles of *Arc* and *Fos* in CA1 samples in IE (red), SC (gray) and EE (blue) conditions. The merged profiles of three independent samples are shown. **D**. PCA of nuRNA-seq profiles of DG neurons in IE, SC an EE conditions (n = 3 per rearing condition). **E**. Heatmap of DEGs in nuRNA-seq profiles of DG neurons in IE, SC and EE conditions (padj < 0.1). The gene expression fold change of each gene in the CA1 SC vs DG SC comparison is also shown. IEGs are highlighted in yellow. **F**. Snapshot of nuRNA-seq genomic profiles of *Arc* and *Nr4a1* genes in DG samples in IE (red), SC (gray) and EE (blue) conditions. The merged profiles of three independent samples are shown. **G**. The effect of EE and IE on the expression levels of the top 200 KA-induced genes in CA1 and DG excitatory neurons is shown. The log_2_FC of all detected genes and a randomly selected subset of 200 genes are also plotted as controls. The horizontal line across the violin plot represents the median. Kruskal-Wallis test followed by Dunn’s multiple comparisons test., **** p < 0.0001, ns not significant.

In DG granule neurons, PCA revealed slight segregation of EE samples, while IE and SC samples were intermingled (**Fig. 4D**). In agreement with this pattern, IE and SC displayed similar expression levels among the 74 DEGs identified (padj < 0.1; **Supp. Table 6** and **Supp. Fig. S6B**), while EE was responsible for most transcriptional changes, leading to the upregulation of the majority of the DEGs (**Fig. 4E**). Almost all DEGs were enriched in CA1 compared to DG, suggesting that EE induces a subset of the CA1 transcriptional program in DG granule neurons (**Fig. 4E**). Of the 74 DEGs identified in DG, only 3 overlapped with the DEGs in CA1, indicating that rearing conditions modulate distinct gene sets in pyramidal and granule neurons. However, remarkably, we retrieved multiple IEGs (*Arc, Nr4a1, Dusp5, Pcsk1, Lbh, 1700016p03rik*) upregulated under the EE condition (**Fig. 4E, F**), demonstrating that EE enhances the activity-driven transcriptional program also in the DG.

Next, to analyze the global behavior of the activity-induced transcriptional program in response to environmental conditions, we examined the EE/SC and IE/SC fold-change of the 200 genes most strongly induced by KA in the hippocampus (*29*). Remarkably, EE led to a significant up-regulation of the activity-induced gene program both in CA1 and in DG, while IE robustly downregulated this set of genes in the CA1 but had no apparent effect in the DG (**Fig. 4G**). In agreement, GSEA analysis of KA-induced genes revealed a strong enrichment of this gene set in EE compared to SC both for CA1 and DG neurons, and an enrichment in SC compared to IE exclusively in CA1 neurons (**Supp. Fig. S6C**). We also compared our nuRNA-seq data with four published transcriptomic profiles of different brain regions in response to EE (**Supp. Fig. S6D**) (*30–33*). Most of the shared genes were IEGs, with *Fosb* being identified as an upregulated DEG in all the studies considered. Collectively, these findings strongly indicate that EE condition stimulates the activity-dependent transcriptional program in hippocampal excitatory neurons.

Altogether, our nuRNA-seq approach revealed that IE and EE exert opposite and cell-type specific effects on the transcriptional profiles of CA1 and DG excitatory neurons. While IE predominantly drives gene repression in CA1 pyramidal neurons, EE primarily promotes gene expression in DG granule neurons and to a lesser degree in CA1 pyramidal neurons, leading to the bidirectional modulation of the activity-dependent genes in both cell types.

### EE increases chromatin accessibility and modulates DNA methylation in a subset of granule neuron enhancers

Regarding chromatin accessibility, our ATAC-seq analysis did not uncover any differentially accessible regions (DARs) among the three housing conditions in the CA1 samples, indicating that exposure to the different environments did not trigger strong changes in the chromatin of CA1 pyramidal neurons (**Supp. Fig. S7A, left**). Conversely, 225 DARs with increased accessibility in EE compared to SC and IE were identified in DG granule neurons (**Fig. 5A, Supp. Fig. S7A**, **right**, and **Supp. Table 7**). This finding is consistent with the evidence that most DEGs in DG neurons were up-regulated under EE (**Fig. 4E**). Notably, 221 out of 225 DARs were classified as enhancers in DG neurons. Since enhancers can regulate gene expression through long range interactions with promoters, this may account for the small overlap (three genes: *Etl4, Ext1* and *Asap1*) between DEGs and the genes closest to the DARs.

**Figure 5.**
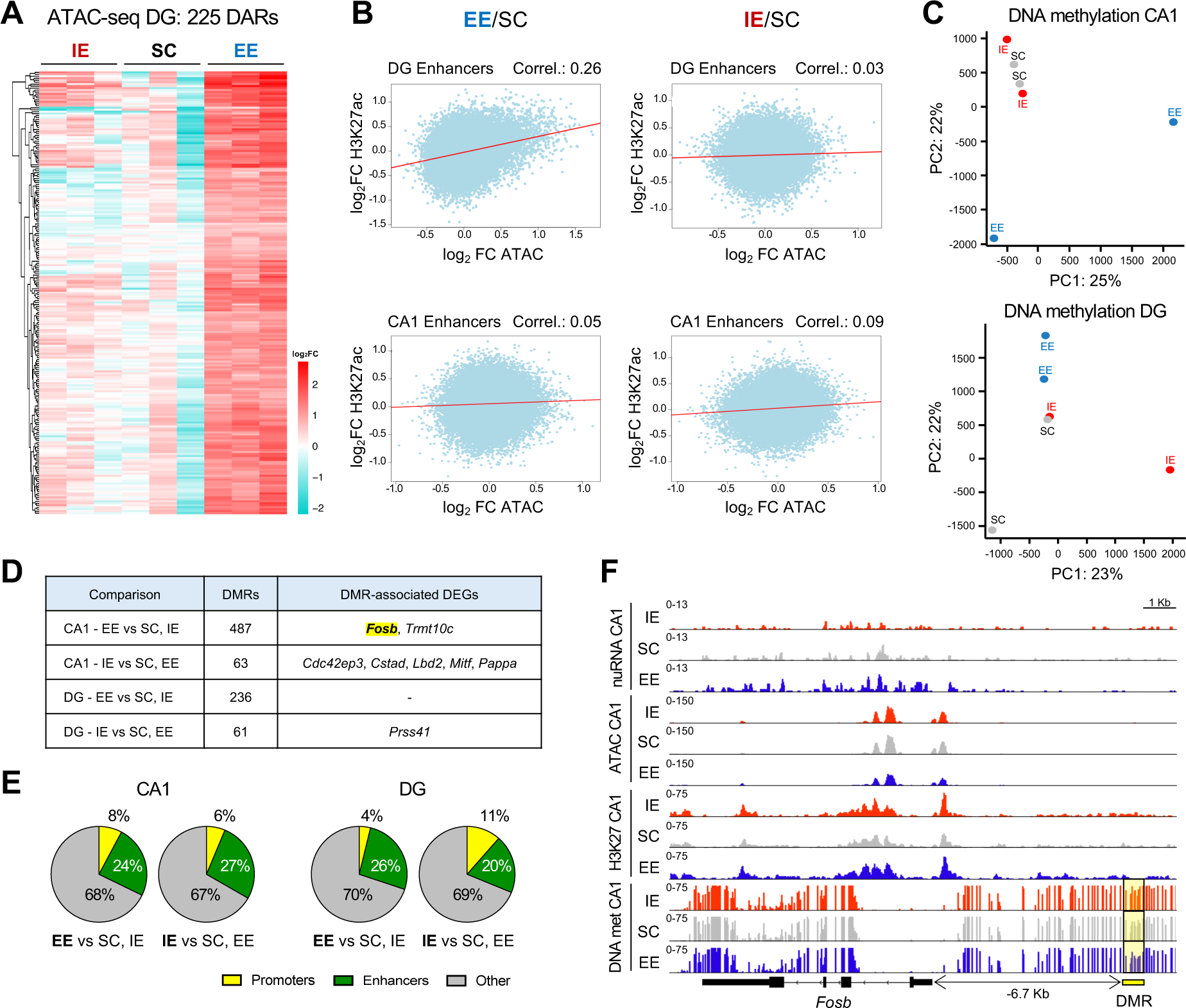
EE modulates chromatin accessibility and DNA methylation in subsets of granule neurons enhancers. **A**. Heatmap of DARs in ATAC-seq profiles of DG SUN1-GFP^+^ neurons in IE, SC and EE conditions (padj < 0.1). N=3 samples per rearing condition. **B**. Pearson correlation between the fold change in chromatin accessibility and the fold change in H3K27ac levels when comparing EE vs SC (left) and IE vs SC (right), within DG (top) and CA1 (bottom) excitatory neurons enhancers. **C**. PCA of DNA methylation profiles of CA1 (left) and DG (right) SUN1-GFP^+^ neurons in IE, SC an EE conditions. N=2 samples per region per rearing condition. **D**. The number of DMRs and the associated DEGs for each comparison are shown. **E**. Pie charts representing the fraction of DMRs overlapping with promoters and enhancers for each comparison. **F**. Snapshot of nuRNA-seq, ATAC-seq, H3K27ac and DNA methylation genomic profiles of *Fosb* gene in CA1 excitatory neurons for IE (red), SC (gray) and EE (blue) conditions. DNA methylation is represented as the percentage of methylation at a given position. The DMR located 6.7 kb upstream of *Fosb* TSS is highlighted with a yellow rectangle. The merged profiles of multiple independent samples are shown.

In term of histone acetylation, our analyses indicate that rearing conditions do not significantly affect H3K27ac profiles in CA1 and DG excitatory neurons. PCA of H3K27ac profiles obtained by CUT&TAG did not reveal a clear separation of the samples according to rearing conditions (**Supp. Fig. S7B**), and peak calling analysis did not identify any differential peak between environmental conditions for either of the hippocampal layers. We however found a positive correlation between the EE-induced changes in chromatin accessibility and H3K27ac levels within DG enhancers, but not when analyzing the IE-induced changes (**Fig. 5B**). This suggests that a gain in H3K27 acetylation accompanies the increase in enhancers accessibility triggered by EE in DG granule neurons.

Finally, PCA of DNA methylation profiles revealed that EE samples separated from IE and SC samples both in CA1 and DG neurons, suggesting that EE might have a more pronounced impact on DNA methylation profiles (**Fig. 5C**). Based on the PCA analysis, we investigated differential methylated regions (DMRs) between EE and the pool of IE and SC samples and identified 487 and 236 DMRs in CA1 and DG, respectively. Only ∼60 DMRs were found when comparing IE against SC and EE samples in CA1 and DG, in agreement with PCA results (**Fig. 5D** and **Supp. Table 8**). Over 30% of the DMRs were found in promoters or enhancers, suggesting that the DNA methylation changes triggered by rearing conditions might affect gene regulation (**Fig. 5E**). However, like for DARs, the overlap between DMR-associated genes and DEGs was restricted to a few genes (**Fig. 5D** and **Supp. Table 8**). Remarkably, among these hits we identified a DMR located 6.7 kb upstream of *Fosb* gene in the chromatin of CA1 neurons, which showed lower DNA methylation levels after EE (**Fig. 5F**). Given the established role of promoter DNA methylation in gene repression, the DMR located in close proximity of *Fosb* TSS may potentially contribute to *Fosb* upregulation in CA1 neurons in response to EE.

In conclusion, our epigenetic study showed that EE increases the accessibility of a restricted subset of enhancers specifically in DG granule neurons and modifies DNA methylation profiles in regulatory regions of CA1 and DG excitatory neurons, including the *Fosb* promoter. In contrast, IE had reduced effects on chromatin occupancy and DNA methylation in both cell types.

### EE and IE bidirectionally modulate AP-1 activity and its downstream program in a cell-type specific manner

We next investigated whether rearing conditions alter chromatin occupancy by TFs in CA1 and DG excitatory neurons through footprinting analysis of our ATAC-seq dataset. Remarkably, in pyramidal neurons our screen revealed a decreased binding of the activity-induced AP-1 transcriptional complex at enhancer regions in the IE condition compared to SC (**Fig. 6A, left,** and **Fig. 6B, top**). Notably, we found 1,676 enhancer regions predicted to be bound by AP-1 exclusively in SC, and not in IE. Since AP-1 regulates the transcription of key synaptic genes required for memory processes (*46, 47*), the reduced binding of AP-1 to CA1 enhancers regions in IE conditions might underlie the cognitive deficits associated with IE rearing. Importantly, we empirically confirmed that AP-1 binds at these genomic sites upon neuronal activation by performing ChIP-seq against FOS in hippocampal chromatin of mice treated with KA (**Fig. 6C**). Additionally, we also detected a decrease in Myocyte-enhancing factor 2 (MEF2) binding at enhancer regions after EE (**Fig. 6A, left**). MEF2 proteins are a family of evolutionarily conserved TFs with a distinctive role as memory constrainers (*48, 49*). Thus, the EE-induced decrease of MEF2 occupancy at CA1 enhancer regions might contribute to the cognitive improvement observed in EE-reared mice.

**Figure 6.**
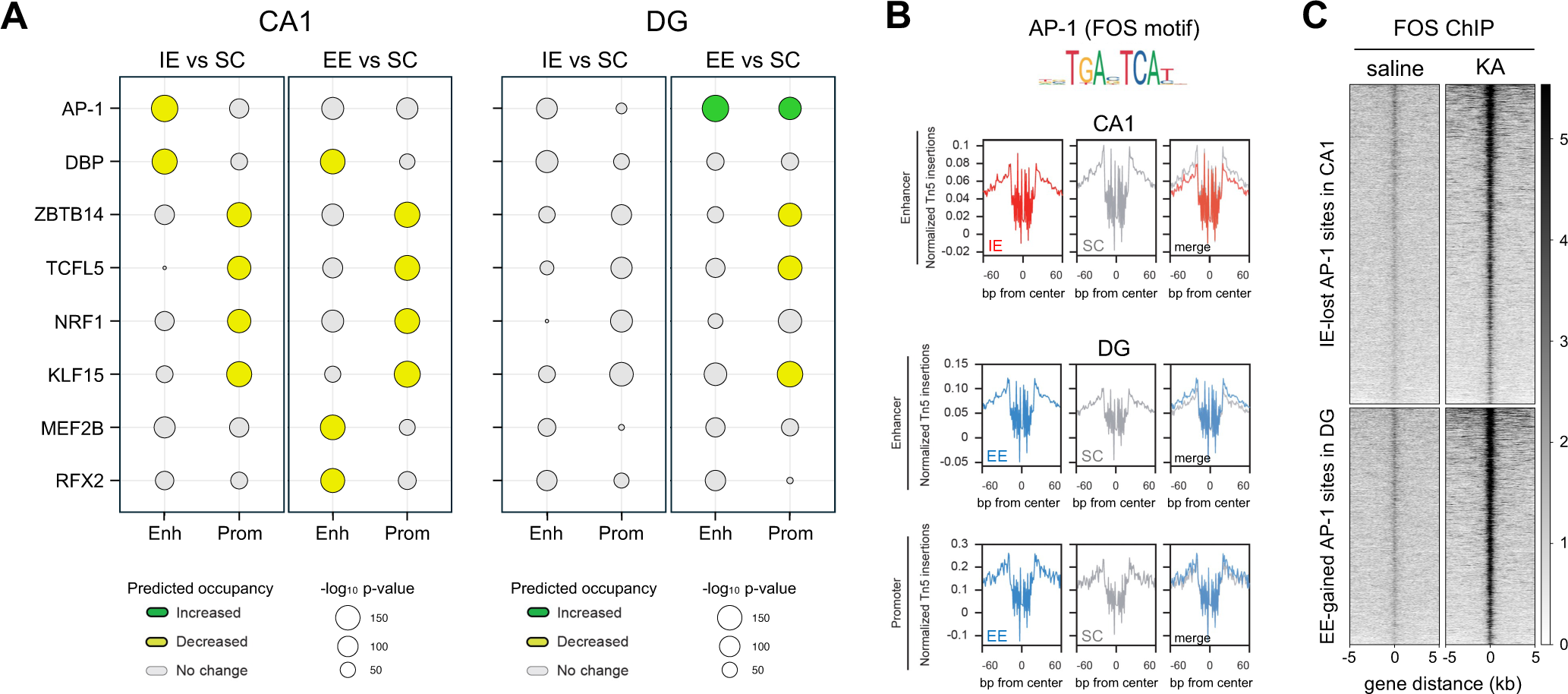
EE and IE modulate AP-1 activity and its downstream transcriptional program in a cell-type specific manner. **A**. TF binding prediction in enhancers and promoters comparing EE vs SC and IE vs SC in CA1 (left) and DG (right) excitatory neurons. The two TFs with the strongest binding change for each comparison are shown (p < 0.05 and binding differential score > 0.25 or < -0.25). Circle size indicates motif enrichment p-value, and circle color refers to the predicted occupancy by the TFs. **B**. Digital footprint for AP-1 (FOS motif) in CA1 neuron enhancers comparing IE and SC (top) and in DG neuron enhancers and promoters (bottom) comparing EE and SC. Values correspond to normalized Tn5 insertions. **C**. FOS ChIP-seq was performed on hippocampal chromatin obtained from mice treated with saline or KA 1h. The heatmaps shows the read density in IE-lost AP-1 sites in CA1 enhancer regions and in EE-gained AP-1 sites in DG enhancer regions.

In granule neurons, footprint analysis revealed an enrichment of AP-1 binding in both promoters and enhancers of DG neurons from EE mice (**Fig. 6A, right,** and **Fig. 6B, bottom**). In particular, 1,242 regions located in DG enhancers are occupied by AP-1 exclusively in the EE condition. FOS ChIP-seq validated these regions as bound by AP-1 after neuronal activation (**Fig. 6C**). Taken together, our footprint analyses indicate that rearing the animals in EE stimulates AP-1 DNA-binding activity in DG granule cells, while IE condition leads to reduced AP-1 signaling in CA1 pyramidal neurons. Given the key role of AP-1 in the transcriptional control of synaptic genes, these results suggest that the beneficial and detrimental effects of EE and IE conditions, respectively, on memory capacities, might rely on the opposite modulation of AP-1 activity by rearing conditions.

### Fos is required for EE-induced cognitive enhancement

To test whether the AP-1 component FOS is required for mediating the environmental effects on cognition, we generated inducible forebrain-specific knockout mice in which *Fos* is conditionally ablated in excitatory neurons (*Fos^f/f^* x *pCamKIIα-CreER^T2^* mice administered with tamoxifen, hereafter referred to as *Fos-ifKO* mice), and tested their performance in the MWM after EE rearing. Previous research has shown that mice lacking neuronal *Fos* can learn to localize the hidden platform in the MWM as efficiently as controls (*50*). Also, multiple laboratories, including ours, have shown that EE improves spatial navigation in the MWM (*5–7, 34*) (**Fig. 1G**). However, the interaction between *Fos* deficiency and EE has not been tackled.

Female *Fos-ifKO* mice were housed in EE conditions during the first 3 months after weaning, before testing their cognitive abilities in the MWM task. Female littermates lacking the *CreER^T2^* transgene, and thus retaining *Fos* expression after tamoxifen injections were also reared in EE cages and served as control. Notably, *Fos-ifKO* mice matched control performance in visible and hidden phases but showed longer latencies in the transfer phase, indicating impaired cognitive flexibility under EE conditions (**Fig. 7A** and **Supp. Fig. S8A, B**). Additionally, *Fos-ifKO* animals showed a modest but significant memory impairment in the NOR assay compared to controls (**Fig. 7B** and **Supp. Fig. S8C**). These results support the conclusion that the AP-1 complex plays a role in mediating the beneficial effect of EE on cognitive abilities.

**Figure 7.**
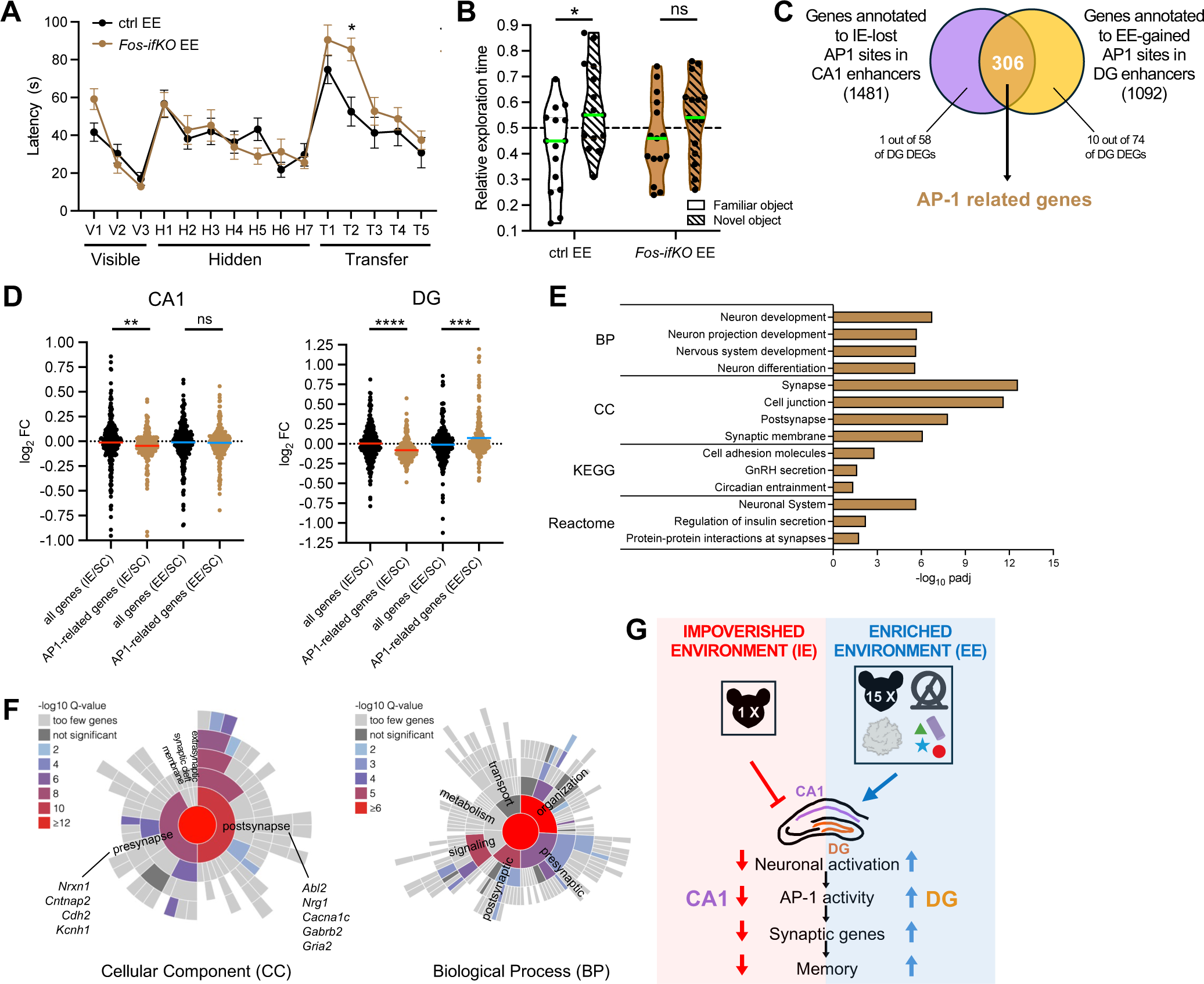
*Fos* is required for EE-induced cognitive enhancement. **A**. Morris Water Maze. The latency to reach the platform was measured for EE-raised ctrl and *Fos ifKO* mice. n = 14 mice per group. Repeated measures ANOVA with correction for multiple comparisons using Šídák’s test. Visible and hidden phase: day ****, genotype ns. Transfer phase: day ****, genotype *. * p < 0.05, ns not significant. **B**. Novel Object Recognition. The relative time spent exploring the familiar and the novel object are shown. The green horizontal line across the violin plot represents the median. n = 15 mice per group. Two-tailed Mann-Whitney test, * p < 0.05, ns not significant. **C**. Venn diagrams showing the overlap between the genes annotated to the IE-lost AP-1 sites in CA1 enhancers (violet), and the genes annotated to the EE-gained AP-1 sites in DG enhancers (orange). **D**. The effect of EE and IE on the expression levels of the 306 AP-1 related genes in CA1 (left) and DG (right) excitatory neurons is shown. The fold change of a same-sized randomly selected subset of genes is plotted as a control. The horizontal line across the violin plot represents the median. Kruskal-Wallis test followed by Dunn’s multiple comparisons test. ** p < 0.01, *** p < 0.001 **** p < 0.0001, ns not significant. **E**. Gene Ontology (GO) enrichment analysis of the 306 AP-1 related genes. The 3-4 significant GO terms with higher -log_10_padj for Biological Process (BP), Cellular Component (CC), KEGG pathway and Reactome pathway are shown. **F**. SynGO analysis of Cellular Component (left) and Biological Process (right) for the 306 AP-1 related genes. Examples of genes which have associated with defects in learning and memory or to ASD (SFARI database) are shown. **G**. Working model. EE and IE bidirectionally modulate the activity-dependent AP-1 pathway in CA1 and DG excitatory neurons in a cell-type specific manner, leading to the transcriptional regulation of AP-1 target genes critical for synaptic function and plasticity, and thus contributing to the opposite effects of EE and IE on memory abilities.

AP-1 contributes to the regulation of a second wave of activity-driven transcription, affecting genes encoding synaptic effector proteins (*44*). To identify the genes which might mediate AP-1 effect on memory capacities, we annotated the AP-1 binding sites with altered AP-1 occupancy in our footprint analysis to their closest gene. The 1,676 CA1 enhancer regions which lose AP-1 binding with IE were annotated to 1,481 genes, and the 1,242 DG enhancers which gain AP-1 occupancy exclusively with EE were annotated to 1,092 genes. By intersecting these two groups of genes, we found 306 genes annotated to enhancers whose AP-1 occupancy is strongly sensitive to environmental conditions, both in CA1 and in DG neurons (**Fig. 7C** and **Supp. Table 9**). Remarkably, the average expression of these 306 AP-1 related genes was decreased by IE both in CA1 and in DG neurons and increased by EE in DG neurons (**Fig. 7D**), suggesting that AP-1 binding at enhancers contributes to transcriptional regulation in response to environmental conditions. Furthermore, the genes annotated to sites gaining EE-induced AP-1 occupancy in DG neurons included 10 out of 74 DG DEGs (hypergeometric p-value = 0.0098), further highlighting the role of AP-1 in regulating gene expression in granule neurons under EE (**Fig. 7C**). GO analysis of the 306 AP-1 related genes revealed significant enrichment in multiple terms related to synapses (**Fig. 7E**), with over one fifth of the genes annotated as synaptic genes in the manually curated SynGO database (**Fig. 7F** and **Supp. Table 9**). Network analysis using STRING (*51*) (enrichment p-value < 10^-16^) indicated that the proteins coded by AP-1 related genes exhibited physical or functional interactions significantly higher than chance, supporting the hypothesis that they participate in common synaptic processes.

Genetic alterations in numerous AP-1 regulated genes have been associated with defects in learning and memory in both humans and animal studies, such as *Csmd1* (*52*), *Cacna1c* (*53*), *Cntnap2* (*54*), *Nrxn1* (*55*), *Abl2* (*56*), *Nrg1* (*57*), and *Mef2c* (*58, 59*). Remarkably, according to the SFARI database (*60*), 64 of the 306 AP-1 related genes have been associated to ASD (**Supp. Table 9**), suggesting that rearing conditions may influence the expression of genes critical for intellectual and social abilities through AP-1 signaling.

These findings underscore the critical role of AP-1 bridging environmental influences during the juvenile period to transcriptional regulation of synaptic genes and highlight potential therapeutic targets for enhancing cognitive function and addressing neurodevelopmental disorders.

## Discussion

Our experiments indicate that rearing mice in EE or IE during the juvenile period has enduring effects on cognitive capacities. Several weeks after being returned to standard cages, EE-SC mice exhibited improved associative learning and spatial memory. Conversely, IE-SC mice performed worse than control littermates in several tasks but showed overall a more limited and task-specific impact on cognitive functions. What underlies these long-lasting effects at the molecular level? In this study, we purified the nuclei of hippocampal excitatory neurons from female mice housed in EE, IE, and standard conditions and profiled their transcriptional and epigenetic signatures to identify molecular changes underlying the effects of early post-weaning environmental conditions on cognitive functions. Our multiomic analysis identified a set of synaptic genes regulated by the transcription factor AP-1 in CA1 and DG excitatory neurons in response to housing conditions. Notably, EE enhanced, whereas IE suppressed, neuronal activation and downstream AP-1 pathway activity in a cell-type-specific manner. This differential regulation of AP-1 target genes, critical for synaptic function and plasticity, contributes to the opposing effects of EE and IE on cognitive abilities (**Fig. 7G**).

### Cell type-specific transcriptional and chromatin signatures of rearing conditions

Comparison of CA1 and DG samples from control mice revealed significant and correlated differences in gene expression, chromatin accessibility, histone acetylation, and DNA methylation between these two neuronal types. The distinct chromatin profiles confer each neuronal type with a unique set of enhancers, which shapes its specific transcriptome. These datasets serve as a valuable resource for the neuroscience community, enabling the computational deconvolution of whole-hippocampus genomic data.

Notably, CA1 neurons exhibited higher expression of activity-dependent transcriptional programs compared to DG neurons. This aligns with observed differences in the basal expression of immediate early genes (IEGs) such as *Arc* and *Fos* across hippocampal layers (**Fig. 3F**, see also (*61, 62*)), reflecting the higher tonic activity of CA1 neurons relative to sparsely active DG neurons. These differences in gene expression likely contribute to the distinct activation dynamics, plasticity mechanisms, and mnemonic functions of pyramidal and granule cells. CA1 pyramidal neurons are highly excitable, with lower action potential thresholds, higher firing rates, and a well-defined afterhyperpolarization (AHP) that supports spike-frequency adaptation. In contrast, DG granule neurons, which exhibit higher input resistance, lower excitability, and require stronger inputs to reach firing thresholds (*35, 39, 63*), generate sparse and temporally precise spiking that is critical for pattern separation and filtering noisy inputs (*64, 65*). These differences underscore the complementary roles of CA1 and DG neurons in hippocampal circuits, with environmental conditions during rearing further amplifying these distinguishing features.

Gene sets regulated by EE and IE were largely distinct in CA1 and DG neurons. DG granule neurons responded primarily to EE, which promoted gene expression, while CA1 pyramidal neurons were more sensitive to IE, which induced gene repression. Moreover, our nuRNA-seq screen revealed bidirectional modulation of activity-dependent gene programs: EE enhanced IEG expression in DG (and to a lesser extent in CA1), whereas IE suppressed IEG expression in CA1. Chromatin configuration also differed between the conditions. EE increased chromatin accessibility in a subset of DG enhancers, while IE had negligible effects. Additionally, EE induced hundreds of DMRs in both CA1 and DG neurons, with over 30% located in promoters or enhancers, suggesting roles in gene regulation and aligning with previous reports of EE-driven DNA methylation changes in hippocampal neuronal genes (*66*). However, we observed limited overlap between DEGs and genes near DARs or DMRs. This could stem from enhancers regulating distant genes or from chromatin and methylation changes priming regulatory regions without immediately affecting basal gene expression.

### Changes in AP-1 activity reflect neuroadaptation to rearing conditions

Our cell-type specific multiomic analysis identified only a few hundred genes and regulatory regions affected by housing conditions during the juvenile period, consistent with previous studies showing that EE impacts specific loci (*32, 33*). Whereas the separation of pyramidal and granule neurons has likely increased the resolution of our screening compared to previous studies, greater inter-individual variability induced by EE (*11, 67*) and the occurrence of environment-driven changes in discrete neuronal ensembles within the hippocampus may have avoided the detection of subtler differences.

Nevertheless, our analysis identified the activity-dependent TF AP-1 and its downstream transcriptional network as the primary signaling axis responding to rearing conditions in hippocampal neurons. Multiple findings support this conclusion. First, TF footprint analysis showed that EE increases AP-1 binding at regulatory regions in the DG, while IE reduces it in CA1 enhancers. Sparse *Fos* expression in DG granule cells may enhance the detection of EE-induced AP-1 activation, whereas weak basal *Fos* expression in CA1 pyramidal neurons facilitates detecting reduced AP-1 activity with IE. Second, over 300 genes associated with AP-1-regulated enhancers were identified, with their collective expression increased under EE and decreased under IE. These genes are enriched in synaptic functions, suggesting that AP-1 modulates their levels to support synaptic changes driven by rearing conditions. Third, both *Fos* and *Fosb*, key AP-1 components, were significantly regulated by environmental conditions in CA1 pyramidal neurons, in line with previous reports (*30, 68*). A DMR in the proximity of *Fosb* TSS showed reduced DNA methylation in EE-raised mice, suggesting a chromatin mechanism contributing to *Fosb* upregulation, consistent with evidence linking DNA demethylation to the expression of activity-regulated genes (*69*). Our findings thereby position AP-1 as a key driver of hippocampal circuit adaptation to EE and IE, directly impacting memory performance. Different subunits can be part of the AP-1 dimeric complex (JUN, JUNB, JUND, FOS, FOSB, FOSL1, FOSL2) (*70*). Remarkably, and despite the documented redundancy among AP-1 components (*71, 72*), deleting *Fos* in excitatory neurons impaired memory performance under EE, underscoring AP-1 role in orchestrating memory-related gene expression (*46*). Future studies should also explore the impact of *Fosb* ablation on EE-induced effects since this subunit emerged in our screen as the most sensitive AP-1 component responding to EE. This finding is consistent with earlier observations on this TF (*31–33*). For instance, the IEG ΔFosB, a truncated product of the *Fosb* gene, is induced in the hippocampus by environmental novelty and long-term exercise, modulates spine density in the CA1, and is required for hippocampal-dependent learning and memory (*73, 74*). Unlike other proteins in the FOS family that are rapidly but transiently induced in response to various stimuli, ΔFosB gradually accumulates during stimulation and remains in the brain for weeks due to its exceptional stability (*75, 76*). This property may allow ΔFosB to drive transcriptional changes that persist well beyond the removal of the initial stimulus, making it an ideal candidate to mediate the long-term consequences of rearing conditions on cognition.

Altogether, our multiomic approach revealed that rearing conditions modify the expression levels, DNA-binding activity, upstream chromatin regulatory mechanisms, and downstream transcriptional effects of AP-1. By demonstrating the lasting benefits of EE on memory, our findings support the translational potential of interventions combining physical exercise, cognitive stimulation, and social interaction as non-pharmacological therapies to counter cognitive impairment or decline (*20, 22*). Manipulating AP1 activity also offers therapeutic potential, enabling modulation of downstream pathways and target genes simultaneously with potential far-reaching implication for the treatment of human cognitive disorders.

## Methods

### Mouse strains and treatments

Mice were kept in a controlled environment with a constant temperature (23 °C) and humidity (40-60%), on 12 h light/dark cycles, with food and water *ad libitum*. They were maintained in a sterile room located within the Animal House at the Instituto de Neurociencias (CSIC-UMH). All animal experiments were performed in agreement with Spanish and European regulations and received approval from the Institutional Animal Care and Use Committee. All experiments were performed on female mice due to frequent fights between males in EE conditions and the difficulty of regrouping male mice after EE or IE. For EE, large cohorts of 10-15 3-week-old C57BL6/J female mice were housed together for 3 months in an ample cage (60 cm x 140 cm x 22 cm) with nesting material, wheels, tunnels and various toys of different shape and texture, which were changed every two weeks. For EI, 3-week-old C57BL6/J female mice were housed individually for 3 months in a small cage (10 cm x 20 cm x 12 cm) without nesting material and placed inside a special armchair, shielding them from social, acoustic and visual stimuli. Female mice kept in standard cages (SC) were raised in group of 4-5 mice per cage, with nesting material. The *pCamKIIα-CreER^T2^* mouse strain (*77*) was crossed with the *pCAG-[STOP]-Sun1-GFP* mouse line (*78, 79*) (stock #021039 from Jackson Laboratory) to label the nuclei of forebrain principal neurons with the nuclear envelope protein SUN1 fused with the reporter GFP, in a tamoxifen-dependent manner (*pCamKIIα-CreER^T2^* x *pCAG-[STOP]-Sun1-GFP*). To remove the STOP cassette and enable *Sun1-GFP* expression, *pCamKIIα-CreER^T2^* x *pCAG-[STOP]-Sun1-GFP* mice received 5 intragastrical administrations of tamoxifen (Sigma Aldrich, 20 mg/mL dissolved in corn oil) orally every other day. To delete *Fos* gene in forebrain excitatory neurons, a mouse carrying floxed *Fos* alleles (*f/f-Fos;* stock #037115 from Jackson Laboratory) (*80*) was crossed with the *pCamKIIα-CreER^T2^* line to obtain mice homozygotes for the floxed *Fos* allele and carrying one copy of the inducible *CreER^T2^* transgene. 3-week-old female mice were transferred to EE cages for 3 months and received 5 intragastrical administrations of tamoxifen every other day starting from P21. The following primers were used for genotyping: *pCamKIIα-CreER^T2^* (GGTTCTCCGTTTGCACTCAGGA, CTGCATGCACGGGACAGCTCT, GCTTGCAGGTACAGGAGGTAGT), *pCAG-[STOP]-Sun1-GFP* (GCACTTGCTCTCCCAAAGTC, CATAGTCTAACTCGCGACACTG, GTTATGTAACGCGGAACTCC), *Fos^f/f^* (GAAGGGACGCTACTGACTGC, TAGCAGGGGAACAAAGAAGC). For KA experiments, mice were administered with 25 mg/Kg KA (or an equal volume of saline for controls) through intraperitoneal injection 1 h before being euthanized.

### Behavioral tests

All behavioral tests were performed during the light cycle. The mice were allowed to habituate to the behavioral room for at least 45 min before each test. Behavioral equipment was cleaned with 70% ethanol after each test session to avoid olfactory cues. Mice behavior was monitored by the video tracking system SMART (Panlab S.L. Barcelona, Spain).

The Open field test (OF) was conducted in a 40 x 40 x 50 cm arena of white acrylic glass. Mice were placed individually in the centre of the arena and allowed to freely explore it for 10 min. The distance run in the centre and the periphery of the arena, the total distance travelled, and the average speed were analyzed using the Mouse Behavioral Analysis Toolbox (MouBeAT) (*81*) for ImageJ.

The Y-maze was performed in a Y-shaped maze of white transparent Plexiglas. Mice were placed individually in the centre of the maze and allowed to freely explore the arms for 6 min. An entry in an arm occurs when all four limbs of the mouse are within an arm. The sequence of entries in the arms was manually scored. An alternation is defined as three consecutive entries into the three different arms. The alternation percentage was calculated with the following formula: n. of alternations * 100/ n. of entries -2.

For contextual fear conditioning (CFC) protocol, mice were placed in a fear conditioning chamber (Panlab S.L., Barcelona, Spain) equipped with an electrified grid. In the conditioning session, mice were allowed to explore the fear conditioning chamber for 2 min before receiving a single 0.4 mA, 2 s footshock, and then remained in the chamber for 1 additional minute. The time animals remained still (freezing) before and after the footshock was registered through a piezoelectric sensor located at the bottom of the fear box, using the software Pacwin (version 2.0.07, Panlab Harvard Apparatus). 24 h later, for the recall session, mice were returned to the same chamber for 3 min, and their freezing behavior was measured.

For novel object recognition (NOR), habituation and testing took place in 40 x 40 x 50 cm white acrylic glass boxes. In a pilot experiment, mice were exposed for 5 min to different couples of plastic objects to select two that fulfilled the following conditions: mice had to show interest for both objects, with no overt intrinsic preference for one object compared to the other. In the two first days of the test, mice were habituated to the test cage for 5 min. On the training session (day 3), the animals were exposed for 5 min to two identical objects (familiar objects) placed on the opposite sides of the box. 24 h later, for the test session (day 4), the mice were exposed for 5 min to the familiar object and to a novel object. Object exploration was defined as approaching and sniffing the object or touching it with the forepaws. The time spent by the animals in investigating each object was manually scored from video recordings by an observer blind to the animal genotype. Mice with a total exploration time inferior to 5 seconds during the training or the test were excluded from the relative exploration time and discrimination index analyses. The discrimination index was calculated with the formula (T_NOVEL_ - T_FAMILIAR_) / (T_NOVEL_ + T_FAMILIAR_), where T_NEW_ = time exploring the novel object, T_FAMILIAR_ = time exploring the familiar object. Eight-arm radial maze test was performed as previously described (*82*).

Morris water maze test (MWM) was performed within a circular tank with a diameter of 170 cm, which was filled with non-toxic white paint. During the initial three days of the test, referred to as visible phase (V1 - V3), a 10 cm diameter platform with a black flag was provided, and mice were trained to locate it in order to exit the water. From days four to seven, known as the hidden phase (H1 - H4), the platform was submerged beneath the water surface in the centre of the target quadrant, and external cues were placed on the walls of the room. Subsequently, during days eight to twelve, referred to as the transfer phase (T1 - T5), the platform was relocated to a new position, requiring the animals to learn its new location to be able to exit from the water. Each day, every mouse underwent four trials, with inter-trial intervals lasting from 30 to 60 min. The trials continued until the mouse reached the platform or for a maximum of 2 min. If the mice did not find the platform after 2 min, they were gently guided to it. Mice were returned to their cages only after staying on the platform for at least 10 s. 1-minute memory retention probe trials (PT), conducted without the platform, were used to assess memory performance at the end of the hidden (PT2) and transfer phases (PT3).

### CA1 and DG microdissection and Fluorescence-activated nuclei sorting (FANS)

Mice were euthanized via cervical dislocation and the CA1 and DG hippocampal layers were microdissected as previously described (*83*). Briefly, brains were removed from the skull and bisected along the longitudinal fissure of the cerebrum. Then, the olfactory bulbs and cerebellum were removed. Positioning the cerebral hemisphere upwards allowed for the removal of the diencephalon (thalamus and hypothalamus) under a dissection microscope, thereby exposing the medial side of the hippocampus. Hippocampus boundaries were separated from the entorhinal cortex and, subsequently, CA3, CA1 and DG were brought into view. CA3 was initially removed, leaving the remaining DG and CA1 regions to be carefully separated along the septo-temporal axis of the hippocampus. All steps were performed at 4 °C unless indicated otherwise. The accuracy of the dissection was evaluated by RT-qPCR of CA1- and DG-specific markers. Tissues microdissected from 2-3 animals were pooled into a single sample. After tissue dissociation, nuclei were extracted by mechanical homogenization using a 2 ml Dounce homogenizer (Sigma-Aldrich) containing 500 μl of Nuclei Extraction Buffer (NEB; Sucrose 250 mM, Cl 25 mM, MgCl_2_ 5 mM, HEPES-KOH pH 7.8 20 mM, IGEPAL CA-630 0.5 %, spermine 0.2 mM, spermidine 0.5 mM, and 1X proteinase inhibitors (cOmplete EDTA-free, Roche)). Tissues from 3 mice were pooled in each sample, having a total volume of 1.5 ml NEB containing the nuclei for each hippocampal layer. Samples were filtered in a 35 μm mesh capped tube and incubated with 0.01 mM DAPI for 10 min in darkness in a rotator. Nuclei isolation was accomplished by the preparation of a gradient of different densities in which nuclei stay in the interphase. Nuclei were diluted in Optiprep density gradient medium (1114542, Proteogenix) to a final concentration of 22 % Optiprep. For the density gradient, 44 % Optiprep was added to a centrifuge tube followed by 22 % Optiprep containing the nuclei. Additional 22 % Optiprep was added until the tube was filled. After centrifugation at 7500 rpm for 23 min, the phase containing the nuclei was collected in a new tube containing Nuclei Incubation Buffer (NIB: sucrose 340 mM, KCl 25 mM, MgCl_2_ 5 mM, HEPES-KOH pH 7.8 20 mM, spermine 0.2 mM, spermidine 0.5 mM, 1X proteinase inhibitors, newborn calf serum 5 %). Sorting of SUN1-GFP^+^ nuclei was performed in a flow-cytometer FACS Aria III (BD Bioscience) in collaboration with the Omics facility at the Instituto de Neurociencias. Sorted nuclei were pelleted by centrifugation (1,000 g, 7 min, 4 °C) and destined to nuRNA-seq, ATAC-seq, H3K27ac CUT&TAG or DNA methylation assays. For each CA1 and DG sample, 75,000 SUN1-GFP^+^ nuclei were used for ATAC-seq, and underwent tagmentation and DNA purification before storage at -20 °C, whereas the remaining nuclei (approx. 700,000 - 1,000,000) were mixed with TRI-reagent (Merck) and stored at -80 °C for nuRNA-seq. Different groups of mice were used to obtain SUN1-GFP^+^ nuclei for H3K27ac CUT&TAG (150,000 and 200,000 nuclei per CA1 and DG sample, respectively) and WGBS (500,000 and 900,000 nuclei per CA1 and DG sample, respectively).

### RNA extraction, retrotranscription and RT-qPCR

Mice were euthanized via cervical dislocation and the CA1 and DG hippocampal layers were dissected from the brain. All steps were performed at 4 °C unless indicated otherwise. RNA extraction was performed using TRI-reagent (Merck) as recommended by the manufacturer. After treatment with RNase-free DNase I (Qiagen) for 30 min at 25 °C to eliminate genomic DNA, RNA was precipitated with phenol-chloroform-isoamyl alcohol. RNA concentration was measured using NanoDropOne (Themo Scientific). RNA was retrotranscribed to cDNA using the RevertAid First-Strand cDNA Synthesis kit (Thermo Scientific) and analysed by RT-qPCR with PyroTaq EvaGreen (Cultek) in a QuantStudio 3 thermocycler (Applied Biosystems). The following primers were used to evaluate the expression of CA1 and DG markers: *Dsp* (F: GCTGAAGAACACTCTAGCCCA, R: ACTGCTGTTTCCTCTGAGACA), *Tdo2* (F: TTTATGGGCACTCTGCTT, R: GGCTCTGTTTACACCAGTTTGAG), *Ccn3* (F: GTCACCAACAGGAATCGCCAGT, R: TACCTTGTCTGTTACTTCCTC). *Gapdh* was used as a reference gene (F: CATGGACTGTGGTCATGAGCC, R: CTTCACCACCATGGAGAAGGC).

### nuRNA-seq

RNA was extracted from SUN1-GFP^+^ nuclei with TRI-reagent (Merck) as described for the RT-qPCR. rRNA depletion libraries were prepared (TruSeq Stranded Total RNA, Illumina), and single-end 50 bp sequencing was conducted in a HiSeq 2500 apparatus (Illumina). *Fastq* files quality was analysed by FastQC (v0.11.9) (*84*) and adapters were trimmed using TrimGalore (v0.39) (*85*). The reads obtained were aligned to the GRCm38.100 mouse genome (mm10) using STAR (v2.7.9a) (*86*). Mitochondrial reads and reads with mapq < 30 were eliminated using Samtools (v1.13) (*87*). The data analysis was conducted with custom R scripts (v4.1.0, 2021), Rsubreads (v2.6.4) (*88*) and Mus_musculus.GRCm38.100.gtf annotation data. For differential expression analysis and samples normalization, DESeq2 (v1.32.0) (*89*) was used. To generate BigWigs, Deeptools (v3.5.1) (*90*) was used. nuRNA-seq reads in vehicle and after 1 hour of KA injection were aligned to this new reference genome using STAR (v2.6.1a). Reads were then filtered for mapq > 30 using Samtools (v1.9), then counts were calculated with Rsubread (v2.4.3) for the reference Mus Musculus.GRCm38.99.gtf where the sfGFP was included. The differential expression analysis was performed using DESeq2 (v1.30.1). For heatmaps, the log_2_FC for each replicate was calculated over the mean of the normalized counts of the SC replicates. GO analysis was performed with gProfiler (*91*) and SynGO (*92*). For Gene Set Enrichment Analysis (GSEA) (Reimand et al., Nature Protocols, 2018), the complete gene list detected by nuRNA-seq in CA1 or DG was ranked by log_2_FC * -log_10_p-value. As a gene set, the 200 genes most strongly up-regulated in the hippocampus after kainate treatment were used (*29*). Protein network analysis was performed with STRING (*51*) taking into account textmining, experiments and databases as interaction sources. gProfiler, SynGO and STRING analysis were performed after conversion of the mouse genes to their human homologues using SynGO ID conversion tool. For the evaluation of *GFP* expression in KA-treated mice, all the nuRNAseq dataset from (*29*) were used to obtain the *sfGFP* sequence using velvet (v1.2.10) (*93*) and it was introduced as an additional chromosome into the mouse reference genome mm10.

### ATAC-seq

ATAC-seq was conducted following the protocol described in (*94, 95*). Briefly, 75,000 SUN1-GFP^+^ sorted neuronal nuclei were subjected to centrifugation, resuspended in the transposase reaction mixture (TD buffer and Tn5 transposase, Illumina) and incubated at 37 °C for 30 min. DNA extraction was immediately performed (MinElute PCR Purification Kit, Qiagen) and samples were stored at -20 °C. Once all samples had been collected, DNA libraries were generated using Custom Nextera PCR primers. The saturation level of the resulting libraries was monitored using RT-qPCR. DNA libraries were purified by double-sided bead purification using AMPure XP beads (Beckman Coulter) to eliminate primer dimers and fragments exceeding 1,000 bp. First, the sample was mixed with 0.5 X volume of AMPure XP beads, incubated for 10 min, placed on a magnetic rack for 5 min, and the supernatant transferred to a fresh tube. Subsequently, 1.3 X original volume of AMPure XP beads were added to the supernatant, mixed thoroughly, incubated for 10 min and placed on a magnetic rack for 5 min. The supernatant was discarded, and the beads were washed with 80 % ethanol. Libraries were eluted in 20 μl of miliQ H_2_O. Purified libraries were stored at -20 °C.

For ATAC-seq analysis, paired-end, 50 bp length sequencing was conducted in a HiSeq 2500 sequencer (Illumina). *Fastq* files quality was analysed by FastQC (v0.11.9) and TrimGalore (v0.39) was used to trim the adapters. Reads obtained were aligned to the GRCm38.100 mouse genome (mm10) using Bowtie2 (v 2.4.2) (*96*) and filtered to eliminate PCR duplicates using Picard (v2.26.2). Mitochondrial reads and reads with mapq < 30 were eliminated using Samtools (v1.13). Peakcalling was conducted using MACS2 (v2.2.7.1) (*97*), and Diffbind (v3.0.15) (*98*) was used for principal component analysis (PCA). Differential accessibility analysis was done using DESeq2 (v1.32.0). To generate BigWigs, Deeptools (v3.5.1) was used to normalize reads by reads per genomic content. With GenomicFeatures (v 1.50.4; (*99*)) a TxDb object was created from the gtf file “Mus_musculus.GRCm38.100.gtf” and genes were annotated using ChipPeakAnno (v 3.32.0) (*100*). For heatmaps, the log_2_FC for each replicate was calculated over the mean of the normalized counts of the SC replicates.

For Transcription Factor (TF) binding prediction, accessible regions were classified into promoters-like or enhancers-like regions using bedtools (v2.30.0). Specifically, accessible regions that mapped to annotated mouse promoters (GRCm38) were considered promoter-like, and the rest were considered enhancers-like and tested for different epigenetic marks (H3K27ac, ATAC-seq reads merged from 3 SC replicates, H3K4me1, H3K4me3, CBP and RNA Pol II). These defined promoters and enhancers regions were used for TF footprints analysis using Transcription factor Occupancy prediction By Investigation of ATAC-seq Signal (TOBIAS, v0.12.9) (*101*). Biological replicates for each condition (IE, SC and EE) were merged by SamTools (v1.13) and the resulting BAM files were corrected for insertion bias of the Tn5 transposase using the command ATACorrect. BigWig files were obtained using the command ScoreBigwig and footprinting scores were assigned using the jaspar vertebrate motif database (JASPAR2020_CORE_vertebrates_non-redundant_pfms_jaspar). Differential TF footprinting for each comparison was calculated with BINDetect command. Bubble plots represented increased and decreased predicted occupancy in red and blue, respectively, based on binding differential score (< -0.25 or > 0.25) and p-value (< 0.05).

### H3K27ac CUT&TAG

H3K27ac CUT&TAG on sorted SUN1-GFP^+^ neurons from IE, NC and EE mice was conducted as described in (*38*) using the anti-H3K27ac antibody from Abcam (ab4729). 75 bp paired-end sequencing was performed using a HiSeq 2500 sequencer. For CUT&TAG analysis, *fastq* files were quality checked using FastQC (v0.11.9). All files presented high quality. Paired-end reads were aligned using Bowtie2 (v2.3.4.3) and were filtered to eliminate duplicates using Picard (v2.26.2). Reads with mapq < 2 were eliminated with Samtools (v1.13). Significant peaks of H3K27ac were called from these aligned files using SEACR (Meers et al., 2019) using the relaxed and norm settings. Differential acetylated regions analysis for neuronal type and experimental condition comparisons was performed using DESeq2 (v1.32.0). To generate BigWigs, Deeptools (v3.5.1) was used to normalize reads by reads per kilobase million. With GenomicFeatures (v 1.50.4) (*99*), a TxDb object was created from the gtf file “Mus_musculus.GRCm38.100.gtf” and genes were annotated using ChipPeakAnno (v 3.32.0) (*100*). For correlational analysis (Fig. 5B), count matrices were obtained from ATAC and H3K27ac CUT&TAG signals for enhancer regions of CA1 and DG separately. Regions with less than 100 counts across all samples were discarded. Log_2_FC of accessibility and H3K27ac levels between rearing conditions for each region were obtained with DESeq2, and Pearson correlation between log_2_FC H3K27ac and log_2_FC accessibility was obtained for each comparison.

### Whole Genome Bisulfite Sequencing (WGBS)

Pelleted nuclei of sorted SUN1-GFP^+^ neurons from IE, NC and EE mice were frozen at -80 °C for later processing. Nuclear pellets were resuspended and incubated with lysis buffer (SDS 0.2 %, Tris pH 8.5 100 mM, EDTA 5 mM, NaCl 200 mM) supplemented with proteinase K for 1h at 56 °C, before DNA extraction with phenol-chloroform method. WGBS was performed using the Swift Accel-NGS Methyl-Seq DNA Library preparation protocol (Swift Biosciences, Ann Arbor, MI). Briefly, first bisulfite conversion is performed on fragmented samples using the EpiTect Bisulfite kit (Qiagen). During the Adaptase step, a short polynucleotide tail along with the first truncated adapter is added at the 3’ of the fragmented and single-stranded bisulfite-converted sample. Then, an extension step is performed followed by a ligation step to attach the second truncated adapter. Finally, full-length adapters are incorporated through an Indexing PCR. 150 bp paired-end sequencing was performed. For DNA methylation analysis, reads were filtered and trimmed using Trimmomatic v 0.30 and aligned to the mm10 genome using BWA v 0.7.8. Methylation calling was then performed using methylCtools. The CpG differential methylation analysis was carried out with methylSig v 1.16.0 (Park et al., Bioinformatics, 2014) according to the reference manual. DNA methylation calls were processed with the Bsseq library v 1.40.0 (Hansen et al., 2012). A single Bsseq object was created for all samples from each brain region (CA1 and DG, two objects in total). Loci were filtered to include only those with coverage above 5 and below 500. The data were then binned into 500 bp windows. PCA plots were generated on the pre-processed data using the prcomp R function with the scaling option applied to the ratio of methylation (number of methylated cytosines divided by coverage for each locus). Testing for differentially methylated regions (DMRs) was conducted using the DSS wrapper function in methylSig v 2.53.0 (Park and Wu, Bioinformatics, 2016), with condition used as contrast. EE samples were tested against combined SC and IE samples while IE samples against SC and EE. The DMRs were assigned to genes using GREAT v 4.0.4 (McLean et al., Nat Biotechnol, 2010; Tanigawa, PLoS Comput Biology, 2022) with the two nearest genes within 200 kb association rule settings. BigWig files used for genome browser visualizations and profiles show the ratio of methylated cytosines to coverage, multiplied by 100, at single base pair resolution and without coverage-based filtering.

### Integration of epigenetic marks and nuRNA-seq datasets

The statistical significance of the overlap between the DEGs upregulated in CA1 or DG (nuRNA-seq; CA1 SC vs DG SC comparison) and the genes annotated to regions with increased epigenetic marks (DARs, differential H3K27ac peaks, DMRs; CA1 SC vs DG SC comparison) was calculated using GeneOverlap R package (v.1.34.0) (Bioconductor) (Shen, 2024). For DNA methylation analysis, given the high number of DMRs identified between CA1 SC and DG SC, the majority of genes were associated to DMRs with increased DNA methylation in both regions. When the difference in the number of DMRs with increased methylation associated to a gene between the two regions exceeded 10, the gene was annotated to increased DMR in the first region. Average profiles for each epigenetic mark across the DEGs upregulated in CA1 or DG were generated based on computeMatrix values with plotProfile from Deeptools (v3.5.1). The intersection of genomic intervals was performed with the Galaxy tool bedtools Intersect intervals (Bjoern A. Gruening, 2014).

### ChIP-seq

ChIP experiment was conducted on the whole hippocampus after treating the mice with 1 h kainic acid (KA) treatment. Hippocampal tissues were homogenized with the Dounce homogenizer in combination with ice-cold hypotonic NEB fixing buffer (250 mM sucrose, 25 mM KCl, 5 mM MgCl_2_, 20 mM Hepes-KOH pH 7.8, 0.5 % IGEPAL CA-630, 0.2 mM spermine, 0.5 mM spermidine, proteases inhibitor 1X and PFA 1.1%). The homogenate was incubated for 15 min at RT followed by 0.125 M glycine for 5 min to stop the fixation, and washed in the centrifuge twice with ice-cold NEB buffer without PFA. The nuclei pellet was suspended in 400 μl of RIPA lysis buffer (1 % IGEPAL CA-630, 0.1 % SDS, 0.5 % sodium deoxycholate, PBS 1x; pH 8) prior to its sonication in a Sonifier digital SFX 250 Branson for 9 cycles of 15s on/30 s off. After 15 min of centrifugation at 17000 g, the supernatant containing isolated chromatin was collected. Samples were diluted in 4 ml of ChIP Dilution buffer (0.01 % SDS, 1.1 % Triton X-100, 1.2 mM EDTA, 16.7 mM Tris-HCl, 167 mM NaCl; pH 8) and incubated overnight at 4°C with the α-FOS monoclonal antibody (Thermo scientific T.142.5). The next day the chromatin was incubated 3 hours with 60 μL of slurry Dynabeads coated to Protein G (Invitrogen) at 4 °C. Beads were thoroughly rinsed at 4 °C as follows: two washes in RIPA-150 buffer, three washes in RIPA-500 buffer (50 mM Tris-HCl, 500 mM NaCl, 1 mM, EDTA, 0.1 % SDS, 1 % Triton X-100, 0.1 % sodium deoxycholate; pH 8), two washes in RIPA LiCl buffer (50 mM Tris-HCl, 1 mM EDTA, 1% NP-40, 0.7 %, sodium deosycholate, 500 mM LiCl2), and two final washes in TE buffer (10 mM Tris-HCl, 1 mM EDTA; pH 8.0), 5 min each. Samples were resuspended in 200 μl Elution buffer (1% SDS, 0.1 M NaHCO_3_) and treated with 1 μl of 10 mg/mL RNase A (Fermentas) overnight at 65 °C in agitation for crosslink reversion. Samples were treated with 2 μl of 20 mg/mL Proteinase K (Thermo Scientific) for 5 h at 55 °C in agitation. Last, DNA precipitation was performed by phenol-chloroform-isoamyl alcohol (Sigma-Aldrich). 50 bp single-end sequencing was performed using a HiSeq 2500 sequencer. Fastq files were trimmed using trim_galore (v0.6.4_dev; Felix Krueger, https://www.bioinformatics.babraham.ac.uk/projects/trim_galore/), after which they were aligned to the mm10 genome using Bowtie2 (v2.3.4.3) (Langmead et al., 2012). Files were filtered for mapq > 30 using Samtools^3^ (v1.9) (Li et al., Bioinformatics, 2009). For differential analysis, the DiffBind^4^ package (v3.12) (Ramírez et al., 2016) was used, with parameters set to minOverlap = 2 and summits = FALSE to calculate counts, and minMembers = 2 to perform contrasts. For profile generation, bigwig files were created using deepTools^5^ (v3.5.0) (Ross-Ines et al., 2012) and loaded into the Integrative Genome Viewer^6^ (v2.17.4) (Thorvaldsdóttir et al., 2013).

### Immunohistochemistry

Mice were anesthetized with a mix of ketamine/xylazine and perfused with 4% paraformaldehyde (PFA) in PBS, and brains were dissected. After an overnight post-fixation in 4% PFA, brain coronal sections (50 μm) were obtained at the vibratome, washed in PBS and incubated for 1 h at room temperature with blocking buffer (5 % Normal Calf Serum, 5 % Normal Goat Serum, 5 % BSA, % Triton X-100 in PBS). The sections were then incubated with primary antibodies in blocking buffer overnight at 4 °C. After 3 x 5 min washes in 0.3 % Triton X-100 in PBS (PBT), sections were incubated with the corresponding secondary antibody overnight at 4 °C, in the dark. After 3 x 5 min washes in PBT, sections were counterstained with DAPI (Invitrogen) before mounting on slides with Fluoromont Aqueous Mounting Medium (Merck F4680-25ML). The following primary and secondary antibodies were used: rabbit α-FOS (Thermo Fisher Scientific #MA5-15055 (T.142.5), 1:100), rabbit α-ARC (Synaptic Systems #156-003, 1:200), goat anti-rabbit 594 (Invitrogen, 1:2000). Confocal images were obtained using a vertical Confocal Microscope Leica SPEII with a dry 20 X objective lens (NA 1.4), with a 1024 x 1024 collection box. Images were processed as sum projections with Fiji software (Schindelin et al., Nat. Methods, 2012).

### Statistical methods

All statistical analyses were conducted using GraphPad Prism v10 (GraphPad Software, La Jolla CA), R v4.4.1, and RStudio v2024.04.2+764. The statistical test used for each analysis is specified in the corresponding figure legend. When comparing two groups, two-tailed Mann-Whitney test was used. When comparing three groups, Kruskal-Wallis test followed by Dunn’s multiple comparisons test was performed. To calculate the statistical significance of the overlap between two groups of genes, we used the online tool https://systems.crump.ucla.edu/hypergeometric/. The accompanying statistic is based on exact hypergeometric probability.

### Data availability

The genomic data sets generated in this study can be accessed at the Gene Expression Omnibus (GEO) repository using the corresponding accession numbers. GSE288335: nuRNA-seq, ATAC-seq, and H3K27ac CUT&TAG data. GSE285372: ChIP FOS. WGBS data were uploaded to the European Nucleotide Archive (ENA) repository (accession PRJEB62813; secondary accession ERP147933). In addition, we used several previously published datasets. GSE236182: H3K4m1 and H3K4me3 ChIP-seq from mouse adult hippocampus (*102*). GSE133018: CBP and H3K27ac ChIP-seq from mouse adult hippocampus (*103*). GSE125068: RNA Pol II ChIP-seq and nuRNA-seq from mouse adult hippocampus (*29*).

## Acknowledgments

We thank Anne-Laurence Boutillier, Jose Vicente Sanchez-Mut and Rafael Alcala-Vida for critical reading of the manuscript. We thank the personnel of the Mouse facility and the Microscopy and Omics core services at the Instituto de Neurociencias for their assistance. We also thank the collaboration of A. Medrano-Fernandez and J. Medrano-Relinque in preliminary experiments and R. Olivares and C. Racovac for technical assistance. M.A-N. was recipient of a fellowship from Generalitat Valenciana. M.F-R. was recipient of a FPU fellowship from the Spanish Ministry of Education. I.B-M. and S.N. are recipients of fellowships from the Spanish Ministry of Science and Innovation (MICINN). A.B. research is supported by grants FCAIXA HR22-00394 from Fundación LaCaixa, PID2020-118169RB-I00 and PID2023-148442NB-I00 from AEI co-financed by ERDF, AC22-00030/JPND2022-115 from ISCIII and CIPROM/2023/15 from the Generalitat Valenciana. B.W. is supported by the JPco-fuND2 Programme (2022/04/Y/NZ2/00116) from the National Science Centre, Poland. WGBS experiments were performed in collaboration with the DKFZ sequencing facility and received funding from the European Union’s Horizon 2020 research and innovation programme under grant agreement No 824110 (EASI-Genomics). The Instituto de Neurociencias is a “Centre of Excellence Severo Ochoa” (CEX2021-001165-S).

## Author contributions

Conceptualization, A.B.; Methodology, M.A-N., F.M., M.L.H., I.B-M. and B.dB.; Software, M.F-R., M.M., M.K.,S.N, and B.W.; Investigation, M.A-N., F.M., M.L.H., I.B-M. and B.dB.; Data Curation and Visualization, M.A-N., F.M. and M.F-R.; Writing - Original Draft, F.M., M.A-N. and A.B.; Supervision, B.W. and A.B.; Funding Acquisition, A.B.

## References

1. D. Hebb, American Psychologist. 2, 306–7 (1947).

2. S. M. Ohline, W. C. Abraham, Neuropharmacology. 145, 3–12 (2019).

3. C. Rampon et al., Nat Neurosci. 3, 238–244 (2000).

4. A. van Dellen, C. Blakemore, R. Deacon, D. York, A. J. Hannan, Nature. 404, 721–722 (2000).

5. J. L. Jankowsky et al., J. Neurosci. 25, 5217–5224 (2005).

6. J. P. Lopez-Atalaya et al., The EMBO Journal. 30, 4287–4298 (2011).

7. M. Pons-Espinal, M. Martinez de Lagran, M. Dierssen, Neurobiology of Disease. 60, 18–31 (2013).

8. M. G. Prado Lima et al., Proc Natl Acad Sci U S A. 115, E2403–E2409 (2018).

9. M. Nilsson, E. Perfilieva, U. Johansson, O. Orwar, P. S. Eriksson, J Neurobiol. 39, 569–578 (1999).

10. E. Bruel-Jungerman, S. Laroche, C. Rampon, Eur J Neurosci. 21, 513– 521 (2005).

11. G. Kempermann, Nat Rev Neurosci. 20, 235–245 (2019).

12. M. G. Leggio et al., Behav Brain Res. 163, 78–90 (2005).

13. A. L. Takatsu-Coleman et al., Journal of Psychiatry & Neuroscience : JPN. 38, 259 (2013).

14. A. Ieraci, A. Mallei, M. Popoli, Neural Plast. 2016, 6212983 (2016).

15. R. T. Han et al., Front. Mol. Neurosci. 11 (2018), doi:10.3389/fnmol.2018.00246.

16. L. A. Teather, J. E. Magnusson, C. M. Chow, R. J. Wurtman, Eur J Neurosci. 16, 2405–2415 (2002).

17. A. B. Silva-Gómez, D. Rojas, I. Juárez, G. Flores, Brain Res. 983, 128– 136 (2003).

18. R. I. Melendez, M. Lee Gregory, M. T. Bardo, P. W. Kalivas, Neuropsychopharmacol. 29, 1980–1987 (2004).

19. M. L. Gregory, K. K. Szumlinski, Neuroreport. 19, 239–243 (2008).

20. S. Smallfield, C. Heckenlaible, The American Journal of Occupational Therapy. 71, 7105180010p1-7105180010p9 (2017).

21. M. W. McDonald, K. S. Hayward, I. C. M. Rosbergen, M. S. Jeffers, D. Corbett, Front Behav Neurosci. 12, 135 (2018).

22. N. J. Ball, E. Mercado, I. Orduña, Front Psychol. 10, 466 (2019).

23. M. Leon, C. Woo, Front Behav Neurosci. 12, 155 (2018).

24. A. P. Feinberg, M. D. Fallin, JAMA. 314, 1129–1130 (2015).

25. G. Cavalli, E. Heard, Nature. 571, 489–499 (2019).

26. J. J. Day et al., Nat Neurosci. 16, 1445–1452 (2013).

27. I. B. Zovkic, M. C. Guzman-Karlsson, J. D. Sweatt, Learn Mem. 20, 61– 74 (2013).

28. M. Fuentes-Ramos, Á. Barco, Adv Neurobiol. 38, 111–129 (2024).

29. J. Fernandez-Albert et al., Nat Neurosci. 22, 1718–1730 (2019).

30. T.-Y. Zhang et al., Nat Commun. 9, 298 (2018).

31. S. Espeso-Gil et al., Front Mol Neurosci. 14, 664912 (2021).

32. Z. Wassouf et al., Front Cell Neurosci. 12, 112 (2018).

33. R. F. Pérez et al., Nat Commun. 15, 5829 (2024).

34. A. Fischer, F. Sananbenesi, X. Wang, M. Dobbin, L.-H. Tsai, Nature. 447, 178–182 (2007).

35. W. Deng, M. Mayford, F. H. Gage, Elife. 2, e00312 (2013).

36. T. Hainmueller, M. Bartos, Nat Rev Neurosci. 21, 153–168 (2020).

37. M. S. Cembrowski, L. Wang, K. Sugino, B. C. Shields, N. Spruston, eLife. 5, e14997 (2016).

38. H. S. Kaya-Okur et al., Nat Commun. 10, 1930 (2019).

39. R. A. Nicoll, R. C. Malenka, Nature. 377, 115–118 (1995).

40. P. A. Jones, Nat Rev Genet. 13, 484–492 (2012).

41. W. Xie et al., Cell. 148, 816–831 (2012).

42. L. D. Moore, T. Le, G. Fan, Neuropsychopharmacol. 38, 23–38 (2013).

43. F. Chen et al., J Physiol. 594, 3729–3744 (2016).

44. E.-L. Yap, M. E. Greenberg, Neuron. 100, 330–348 (2018).

45. S. Bäck et al., Molecular Therapy - Methods and Clinical Development. 14, 180–188 (2019).

46. C. M. Alberini, Physiological reviews. 89 (2009), doi:10.1152/physrev.00017.2008.

47. E. Benito, A. Barco, Mol Neurobiol. 51, 1071–1088 (2015).

48. C. J. Cole et al., Nature neuroscience. 15, 1255–1264 (2012).

49. A. J. Rashid, C. J. Cole, S. A. Josselyn, Genes, Brain and Behavior. 13, 118–125 (2014).

50. A. Fleischmann et al., J Neurosci. 23, 9116–9122 (2003).

51. D. Szklarczyk et al., Nucleic Acids Res. 51, D638–D646 (2023).

52. L. Athanasiu et al., Brain, Behavior, and Immunity. 61, 209–216 (2017).

53. D. Cosgrove et al., Neuropsychopharmacology. 42, 2612–2622 (2017).

54. A. R. Rendall, D. T. Truong, R. H. Fitch, Behavioural Brain Research. 303, 201–207 (2016).

55. S. Zhang et al., Experimental Neurology. 344, 113806 (2021).

56. A. González-Martín et al., Prog Neurobiol. 205, 102122 (2021).

57. Y. Lu et al., Neuron. 84, 835–846 (2014).

58. C. J. Cole et al., Nat Neurosci. 15, 1255–1264 (2012).

59. A. J. Rashid, C. J. Cole, S. A. Josselyn, Genes Brain Behav. 13, 118– 125 (2014).

60. B. S. Abrahams et al., Mol Autism. 4, 36 (2013).

61. M. K. Chawla et al., Hippocampus. 15, 579–586 (2005).

62. B. N. Jaeger et al., Nat Commun. 9, 3084 (2018).

63. K. A. Alkadhi, Mol Neurobiol. 56, 6566–6580 (2019).

64. M. W. Jung, B. L. McNaughton, Hippocampus. 3, 165–182 (1993).

65. J. K. Leutgeb, S. Leutgeb, M.-B. Moser, E. I. Moser, Science. 315, 961– 966 (2007).

66. S. Zocher, R. W. Overall, M. Lesche, A. Dahl, G. Kempermann, Nat Commun. 12, 3892 (2021).

67. J. C. Körholz et al., eLife. 7, e35690 (2018).

68. M. Hüttenrauch, G. Salinas, O. Wirths, Front Mol Neurosci. 9, 62 (2016).

69. A. Rudenko et al., Neuron. 79, 1109–1122 (2013).

70. F. Bejjani, E. Evanno, K. Zibara, M. Piechaczyk, I. Jariel-Encontre, Biochimica et Biophysica Acta (BBA) - Reviews on Cancer. 1872, 11–23 (2019).

71. A. Fleischmann et al., Genes Dev. 14, 2695–2700 (2000).

72. W.-M. Wang, S.-Y. Wu, A.-Y. Lee, C.-M. Chiang, Journal of Biological Chemistry. 286, 40974–40986 (2011).

73. T. Nishijima, M. Kawakami, I. Kita, PLOS ONE. 8, e81245 (2013).

74. A. L. Eagle et al., J Neurosci. 35, 13773–13783 (2015).

75. T. L. Carle et al., Eur J Neurosci. 25, 3009–3019 (2007).

76. P. G. Ulery-Reynolds, M. A. Castillo, V. Vialou, S. J. Russo, E. J. Nestler, Neuroscience. 158, 369–372 (2009).

77. G. Erdmann, G. Schütz, S. Berger, BMC neuroscience. 8 (2007), doi:10.1186/1471-2202-8-63.

78. W. E. Allen et al., Science. 357, 1149–1155 (2017).

79. L. A. DeNardo et al., Nature Neuroscience 2019 22:3. 22, 460–469 (2019).

80. J. Zhang et al., Nat Genet. 30, 416–420 (2002).

81. E. Bello-Arroyo et al., Front Behav Neurosci. 12, 201 (2018).

82. H. Koshimizu et al., Front. Behav. Neurosci. 5 (2011), doi:10.3389/fnbeh.2011.00085.

83. H. Hagihara, K. Toyama, N. Yamasaki, T. Miyakawa, JoVE (Journal of Visualized Experiments), e1543 (2009).

84. S. Andrews, FastQC A Quality Control tool for High Throughput Sequence Data. Babraham Bioinformatics (2010), (available at https://www.bioinformatics.babraham.ac.uk/projects/fastqc/).

85. F. Krueger, S. Andrews, (2012).

86. A. Dobin et al., Bioinformatics (Oxford, England). 29, 15–21 (2013).

87. H. Li et al., Bioinformatics. 25, 2078 (2009).

88. Y. Liao, G. K. Smyth, W. Shi, Bioinformatics (Oxford, England). 30, 923–930 (2014).

89. M. I. Love, W. Huber, S. Anders, Genome biology. 15 (2014), doi:10.1186/S13059-014-0550-8.

90. F. Ramírez et al., Nucleic acids research. 44, W160–W165 (2016).

91. L. Kolberg et al., Nucleic Acids Research. 51, W207–W212 (2023).

92. F. Koopmans et al., Neuron. 103, 217-234.e4 (2019).

93. D. R. Zerbino, E. Birney, Genome Research. 18, 821 (2008).

94. J. D. Buenrostro, P. G. Giresi, L. C. Zaba, H. Y. Chang, W. J. Greenleaf, Nature methods. 10, 1213–1218 (2013).

95. J. D. Buenrostro, B. Wu, H. Y. Chang, W. J. Greenleaf, Current protocols in molecular biology / edited by Frederick M. Ausubel … [et al.], in press, doi:10.1002/0471142727.MB2129S109.

96. B. Langmead, S. L. Salzberg, Nature methods. 9, 357–359 (2012).

97. Y. Zhang et al., Genome Biology. 9, 1–9 (2008).

98. R. Stark, G. Brown, 1–73 (2022).

99. M. Lawrence et al., PLOS Computational Biology. 9, e1003118 (2013).

100. L. J. Zhu et al., BMC bioinformatics. 11 (2010), doi:10.1186/1471-2105-11-237.

101. M. Bentsen et al., Nature Communications 2020 11:1. 11, 1–11 (2020).

102. B. Del Blanco et al., Nat Commun. 15, 1781 (2024).

103. M. Lipinski et al., Nat Commun. 11, 2588 (2020).

