## Supplemental materials for "Rearing conditions bidirectionally modulate cognitive abilities and AP-1 signaling in hippocampal neurons in a cell type-specific manner"

**SUPPLEMENTARY FIGURES**

- Supplementary Figures S1-S8 (pages 2-9)
- Supplementary Files:
  - o Table S1 related to Figure 2. Differentially expressed genes in the comparison of CA1 and DG nuRNA-seq samples (see attached Excel file, 1 sheet).
  - o Table S2 related to Figure 2. gProfiler GO analysis of DEGs upregulated in CA1 SC (A) and DG SC (B) (see attached Excel file, 2 sheets).
  - o Table S3 related to Figure 2. Differentially accessible regions in the comparison of CA1 and DG ATAC-seq samples and associated genes (see attached Excel file, 2 sheets).
  - o Table S4 relative to Figure 2. Differentially acetylated regions in the comparison of CA1 and DG H3K27ac CUT&TAG samples and associated genes (see attached Excel file, 2 sheets).
  - o Table S5 related to Figure 2. Differentially methylated regions in the comparison of CA1 and DG WGBS samples and associated genes (see attached Excel file, 2 sheets).
  - o Table S6 related to Figure 4. Differentially expressed genes in the comparison of EE/SC/IE nuRNA-seq samples (see attached Excel file, 2 sheets).
  - o Table S7 relative to Figure 5. Differentially accessible regions in the comparison of EE/SC/IE DG ATAC-seq samples and associated genes (see attached Excel file, 2 sheets).
  - o Table S8 relative to Figure 5. Differentially methylated regions in the comparison of EE/SC/IE WGBS samples and associated genes (see attached Excel file, 5 sheets).
  - o Table S9 related to Figure 7. GO analysis of AP1 related genes (see attached Excel file, 3 sheets).

### SUPPLEMENTARY FIGURES

**Supplementary Figure S1**

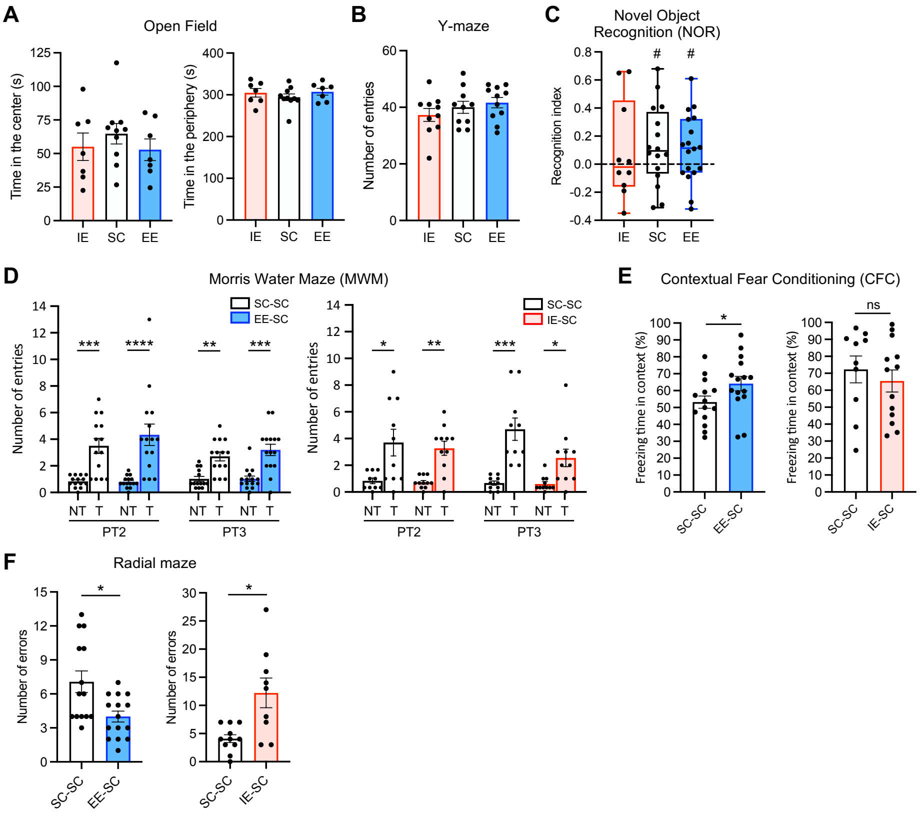


**Supplementary Figure S1 related to Figure 1. Perduring impact of rearing conditions on cognitive capacities. A**. Time spent in the center (left) and in the periphery (right) of the arena in the Open Field test. Mean ± SEM is shown. n = 7-10 mice per group. No statistically significant differences comparing EE vs SC and IE vs SC by Kruskal-Wallis test followed by Dunn's multiple comparisons test. **B**. Number of entries in the arms in the Y-maze. Mean ± SEM is shown. n = 10-11 mice per group. No statistically significant differences comparing EE vs SC and IE vs SC by Kruskal-Wallis test followed by Dunn's multiple comparisons test. **C**. Recognition index in the Novel Object Recognition task. n = 10-18 mice per group. One sample t test against hypothetical value 0, # p < 0.1. No statistically significant differences comparing EE vs SC and IE vs SC by Kruskal-Wallis test followed by Dunn's multiple comparisons test. **D**. Morris Water Maze. Number of entries in the target (T) and non-target (NT) quadrants during the Probe Trials (PT) 2 and 3 for EE-SC vs SC-SC (left) and IE-SC vs SC-SC (right) groups. For NT, the average number of entries in the three non-target quadrant was calculated. n = 10-14 mice per group. Kruskal-Wallis test followed by Dunn's multiple comparisons test for NT vs T comparisons, * p < 0.05, ** p < 0.01, *** p < 0.001, *** p < 0.0001. **E**. Contextual Fear Conditioning. The freezing time in the context session for EE-SC vs SC-SC (left) and IE-SC vs SC-SC (right) groups was measured. Mean ± SEM is shown. n = 10-15 mice per group. Two-tailed Mann-Whitney test, * p < 0.05, ns not significant. **F**. Radial maze. The number of errors as an indicator of working memory impairment was measured for EE-SC vs SC-SC (left) and IE-SC vs SC-SC (right) groups. Mean ± SEM is shown. n = 8-15 mice per group. Two-tailed Mann-Whitney test, * p < 0.05.

**Supplementary Figure S2**

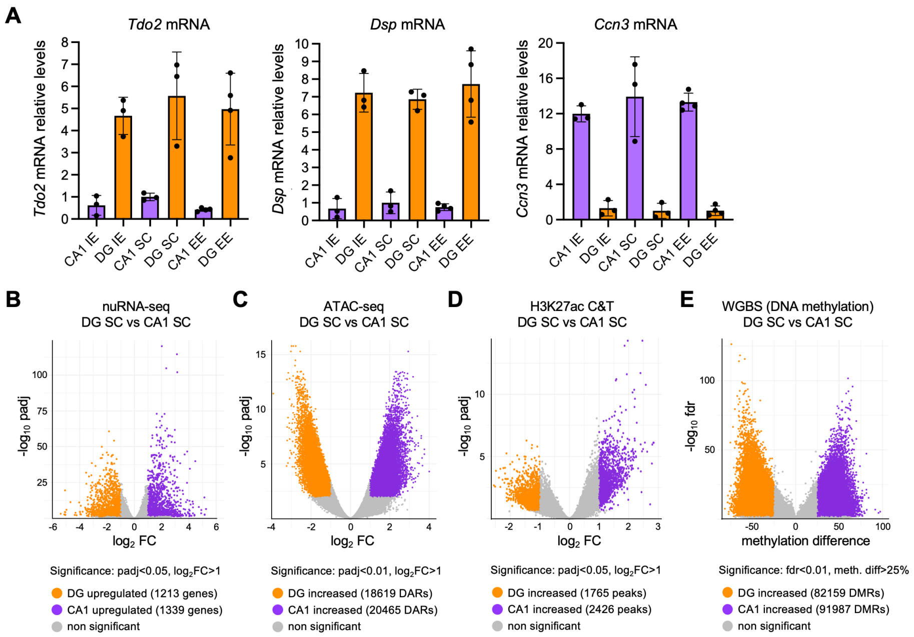


**Supplementary Figure S2 related to Figure 2. Multiomic analysis reveals profound differences in transcriptional and epigenetic profiles of CA1 and DG excitatory neurons. A**. RT-qPCR analysis of *Tdo2* (DG marker; left), *Dsp* (DG marker, middle) and *Ccn3* (CA1 marker, right) mRNA levels in manually dissected CA1 and DG layers from IE, SC and EE mice. Mean ± SEM is shown. n = 3-4 mice per group. Two-tailed Mann-Whitney test comparing all CA1 samples with all DG samples for each gene, $p < 0.0001$. **B-E**. Volcano plot of nuRNA-seq (B), ATAC-seq (C), H3K27ac (D) and DNA methylation (E) differential analysis between CA1 SC and DG SC excitatory neurons. DEGs significantly up-regulated in CA1 and DG neurons are shown in violet and orange, respectively.

**Supplementary Figure S3**

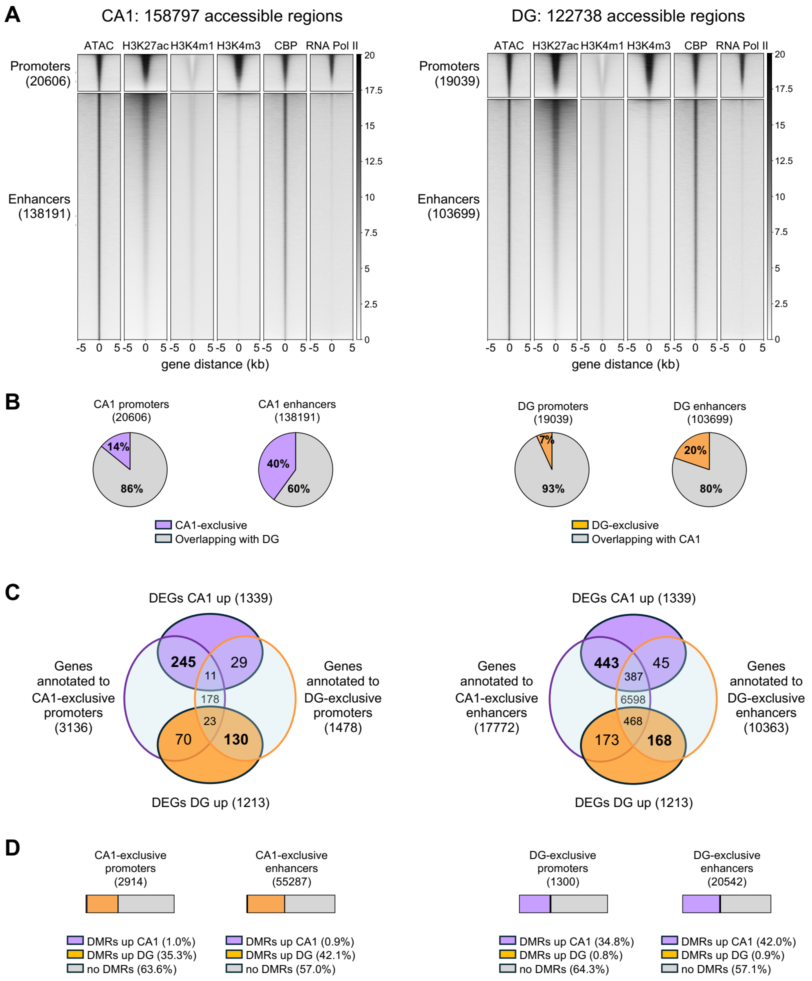


**Supplementary Figure S3 related to Figure 2. Analysis of pyramidal neuron and granule neuron-specific promoters and enhancers. A**. K-mean clustering was applied to identify accessible regions (± 5 Kb window) in the chromatin of CA1 (left) and DG (right) excitatory neurons, using ATAC, H3K27ac, H3K4me1, H3K4me3, CBP and RNAPII binding profiles from hippocampal chromatin. ATAC, H3K27ac and CBP signals denote a regulatory role, while H3K4me3 primarily labels promoter regions. **B**. Fractions of promoters and enhancers exclusive of CA1 (left) or DG (right) excitatory neurons. **C**. Overlap between DEGs upregulated in CA1 or DG neurons and genes annotated to exclusive promoters (left) and enhancers (right) of CA1 and DG neurons **D**. Fractions of increased DMRs within promoters and enhancers exclusive of CA1 (left) and DG (right) excitatory neurons.

**Supplementary Figure S4**

**A**

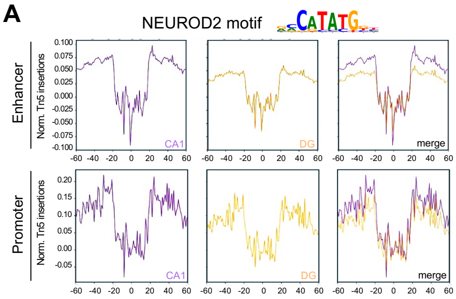


**Supplementary Figure S4 related to Figure 3. Footprint analysis of NEUROD2 motif. A**. Digital footprint for NEUROD2 motif in enhancers (top) and promoters (bottom) comparing CA1 and DG excitatory neurons. Values correspond to normalized Tn5 insertions.

**Supplementary Figure S5**

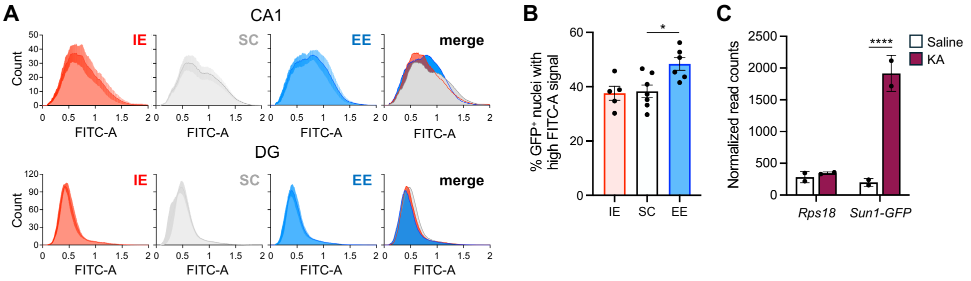


**Supplementary Figure S5 related to Figure 3. EE leads to increased tonic activity in CA1 pyramidal neurons. A**. Flow cytometer histograms (number of events versus FITC-A channel signal, arbitrary units) of SUN1-GFP$^+$ singlet nuclei from CA1 (top) and DG (bottom) excitatory neurons of mice raised in IE, SC and EE conditions. Mean ± SEM of three experiments is shown. **B**. Percentage of CA1 SUN1-GFP$^+$ nuclei with a FITC-A value higher than the established constant gate. Kruskal-Wallis test comparing EE vs SC and IE vs SC, * $p < 0.05$. **C**. Normalized nuRNA-seq *Sun1-GFP* read counts from hippocampi of mice treated with saline or KA and sacrificed 1-hour post-treatment. *Rps18* is used as housekeeping gene. ****, padj = $7.14 \times 10^{-17}$. nuRNA-seq data were obtained from Fernandez-Albert et al., 2019, and analysed with DESeq2.

**Supplementary Figure S6**

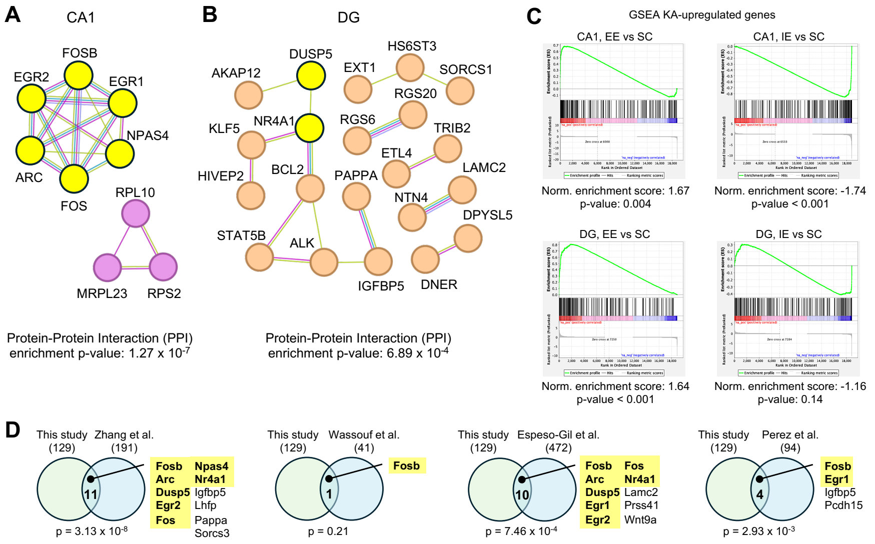


**Supplementary Figure S6 related to Figure 4. Protein network and GSEA analysis of DEGs in CA1 pyramidal neurons and DG granule neurons across environmental conditions. A**, **B**. Network visualization of the protein products of DEGs induced by environmental conditions, in CA1 (A) and DG (B) excitatory neurons. Network analysis was performed using STRING. Only proteins displaying at least one interaction are shown. IEGs are indicated in yellow. **C**. GSEA analysis of the top 200 KA-induced genes in nuRNA-seq when comparing EE vs SC in CA1 (top, left), IE vs SC in CA1 (top, right), EE vs SC in DG (bottom, left) and IE vs SC in DG (bottom, right). **D**. Venn diagrams showing the intersection between the rearing condition-dependent DEGs identified in this analysis and the EE-dependent DEGs found in previous studies. The overlapping genes and the hypergeometric probability are shown. IEGs are highlighted in yellow.

**Supplementary Figure S7**

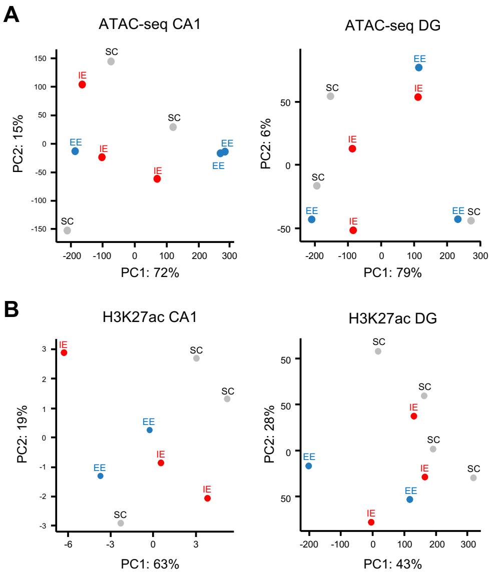


**Supplementary Figure S7 related to Figure 5. PCA of ATAC and H3K27ac profiles in CA1 and DG excitatory neurons across environmental conditions. A**. PCA of ATAC profiles of CA1 (left) and DG (right) SUN1-GFP$^{+}$ neurons in IE (red), SC (grey) an EE (blue) conditions. n = 3 samples per region per rearing condition. **B**. PCA of H3K27ac profiles of CA1 (left) and DG (right) SUN1-GFP$^{+}$ neurons in IE (red), SC (grey) and EE (blue) conditions. n = 2-4 samples per region per rearing condition.

**Supplementary Figure S8**

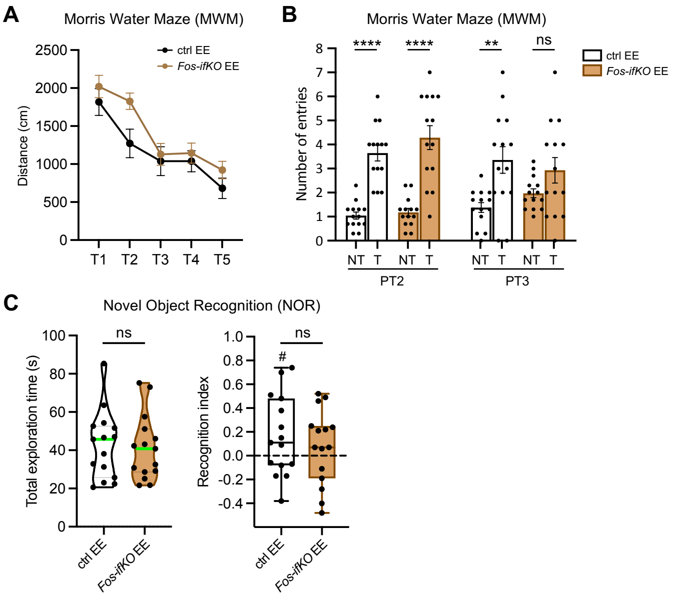


**Supplementary Figure S8 related to Figure 7. Behavioral analysis of *Fos-ifKO* mice. A**. Morris Water Maze. The distance traveled to reach the platform during the transfer phase was measured for EE-raised ctrl and *Fos-ifKO* mice. n = 14 mice per group. Repeated measures ANOVA with correction for multiple comparisons using Šídák's test. Time ****, genotype *. **B**. Morris Water Maze. Number of entries in the target (T) and non-target (NT) annulus during the Probe Trials (PT) 2 and 3 for EE-raised ctrl and *Fos-ifKO* mice. For NT, the average number of entries in the three non-target annulus was calculated. n = 14 mice per group. Kruskal-Wallis test followed by Dunn’s multiple comparisons test for NT vs T comparisons, ** $p < 0.01$, **** $p < 0.0001$, ns not significant. **C**. Novel Object Recognition. The total time spent exploring the identical objects during the training (left) and the recognition index for the test session (right) are shown for EE-raised ctrl and *Fos-ifKO* mice. The green horizontal line across the violin plot represents the median. n = 15 mice per group. Two-tailed Mann-Whitney test, ns not significant. Recognition index, one sample t test against hypothetical value 0, # $p < 0.1$.
